# Rewiring of the epigenome and chromatin architecture by retinoic acid signaling during zebrafish embryonic development

**DOI:** 10.1101/2023.06.13.544553

**Authors:** Marta Moreno-Oñate, Lourdes Gallardo-Fuentes, Pedro M. Martínez-García, Silvia Naranjo, Sandra Jiménez-Gancedo, José L. Gómez-Skarmeta, Juan J. Tena, José M. Santos-Pereira

## Abstract

**Background:** Retinoic acid (RA) functions as a ligand for the nuclear RA receptors (RARs), which regulate the expression of target genes by binding to RA response elements. RA signaling is required for multiple processes during chordate embryonic development, such as body axis extension, hindbrain antero-posterior patterning and forelimb bud initiation. Although some RA target genes have been identified, little is known about the genome-wide effects of RA signaling during *in vivo* embryonic development.

**Results:** Here we stimulate the RA pathway during development by treating zebrafish embryos with all-trans-RA (atRA), the most abundant form of RA, and use a combination of RNA-seq, ATAC-seq, ChIP-seq and HiChIP to gain insight into the molecular mechanisms by which RA signaling control target gene expression. We find that RA signaling is involved in anterior/posterior patterning and development of the central nervous system, participating in the transition from pluripotency to differentiation. atRA treatment also induces alterations in chromatin accessibility during early development and promotes chromatin binding of RARαa and the RA targets Hoxb1b, Meis2b and Sox3, which cooperate in central nervous system development. Finally, we show that RA induces a rewiring of chromatin architecture, with alterations in chromatin 3D interactions that are consistent with target gene expression. This is illustrated by the specific induction of anterior HoxB genes by RARs, among other examples.

**Conclusions:** Altogether, our findings identify genome-wide targets of RA signaling during embryonic development and provide a molecular mechanism by which developmental signaling pathways regulate the expression of target genes by altering chromatin topology.

## INTRODUCTION

Retinoic Acid (RA) is an active metabolite derived from retinol, also known as vitamin A, that regulates multiple developmental processes in chordate animals acting as a diffusible signaling molecule (1). Several isomeric forms of RA exist in cells, including 9-*cis*-RA, 13-*cis*-RA and all-*trans*-RA (atRA), being atRA the major physiological form. RA is produced in two steps by retinol dehydrogenase-10 (Rdh10), which produces retinaldehyde from retinol, and retinaldehyde dehydrogenases (Aldh1a1-3), which convert retinaldehyde to RA (2, 3, 4). On the other hand, RA synthesis is counteracted by dehydrogenase/reductase 3 (Dhrs3), which converts retinaldehyde back to retinol (5). Finally, RA is degraded by cytochrome P450 family enzymes, including Cyp26a1, Cyp26b1 and Cyp26c1 (6, 7). RA is the ligand of RA receptors (RARs), which are transcription factors (TFs) of the nuclear receptor (NR) superfamily that bind chromatin at RA response elements (RAREs) (8, 9). RAREs located nearby RA target genes typically consist of hexameric direct repeats with an interspacing of 2 or 5 bp (DR2 or DR5, respectively) (10), where RARs bind forming a heterodimer with retinoid X receptors (RXRs) to activate target gene expression. There are three RAR isoforms in mammals, RARα, β and γ, as well as three RXR isoforms, RXRα, β and γ, while zebrafish lacks RARβ and have two paralogs of all the other isoforms with different expression patterns (11).

RARs can regulate transcription of important developmental genes in response to RA signaling, being involved in the development of many organs and tissues that include body axis, hindbrain, heart, forelimbs, eyes and reproductive tract (1). During early vertebrate development, RA is produced in the trunk presomitic mesoderm by expression of *Rdh10* and *Aldh1a2*, generating a two-tailed gradient that diffuses anteriorly until the hindbrain and heart, and posteriorly until the caudal progenitor zone (12, 13). At these regions, opposite gradients of Fgf8 signaling and *Cyp26a1* expression limit RA action (14, 15, 16), completing a regulatory mechanism that controls the extension of body axis in vertebrates, although the requirement of RA for this process in zebrafish remains controversial (17). An opposite antagonist interaction between RA and Fgf8 signaling also occurs in heart antero-posterior patterning and in the initiation of forelimbs and fin buds (18, 19).

RA signaling plays a key role in the patterning of the posterior central nervous system, including the hindbrain and the spinal cord (20, 21), and RA treatment induces differentiation of embryonic stem cells to neuroectodermal fates (22). In this regard, *Hox* genes, which encode TFs that are involved in establishing the antero-posterior body axis in animal embryos, as well as other axial structures including limbs and genitalia (23), are well-known downstream targets of RA signaling (24). *Hox* genes respond to RA treatment both in cultured cells and embryos (25, 26), and are organized in several clusters (four in mammals: HoxA, HoxB, HoxC and HoxD; seven in zebrafish) where their spatial organization correlates with their temporal and spatial expression patterns, a property known as collinearity (27). This phenomenon also applies for the relative response of *Hox* genes to signaling pathways as RA, since 3’ genes are more responsive to RA than 5’ genes (28). On the other hand, the binding specificity of Hox TFs to chromatin may be altered by their co-binding with co-factors, including the three amino acid loop extension (TALE) class of homeodomain-containing TFs, such as Pbx, Meis and Prep families (29). In this sense, RA signaling also promotes the expression of *Meis* genes during mouse limb induction in the proximal domains (30), and Meis, Pbx and Hox TFs are known to cooperate in the specification of the hindbrain in *Xenopus* and zebrafish (31).

Despite the large amount of evidence establishing the function of RA signaling and RARs in the regulation of target gene expression, very little is known about their global effect on the chromatin landscape. In this sense, recent findings in mouse and zebrafish pancreas suggest that RA signaling may rewire the chromatin landscape leading to *cis*-regulatory element (CRE) activation (32, 33). Previous studies indicate that active enhancers are engaged in chromatin 3D interactions with target promoters, being established either before or concomitant with gene activation, depending on the context (34, 35). While poised enhancers are already interacting with target promoters in mouse embryonic stem cells (mESCs) (36), enhancer-promoter interactions involving lineage-specific genes are established during neural or erythroid differentiation (37, 38). In the case of RA-induced differentiation, changes in interactions between the RA target gene *Hoxa1* and three nearby enhancers have been recently reported in mESCs (39). However, the effects of RA signaling on global chromatin 3D interactions between enhancers and promoters in an *in vivo* context have not been addressed before.

In this work, we have analyzed the effects of RA treatment during early development in zebrafish embryos at the transcriptomic, epigenomic and chromatin conformation levels, by integrating RNA-seq, ATAC-seq, ChIP-seq and HiChIP. First, we describe the dynamic chromatin binding of one of the zebrafish RARs, RARαa, during gastrulation, segmentation and phylotypic stages. Then, we treat embryos with RA and analyze the global effects at the transcriptomic and epigenomic levels using RNA-seq and ATAC-seq, respectively, uncovering thousands of genes and putative CREs responding to RA signaling. Leveraging ChIP-seq experiments, we show that RA treatment induces chromatin binding not only of RARαa, but also of Hoxb1b, Meis2b and Sox3 TFs, which cooperate to promote the development and patterning of the central nervous system. Finally, we assess chromatin 3D interactions altered by RA signaling using HiChIP of active promoters and find a link between miss-regulation of gene expression and changes in promoter contacts that is illustrated by bona fide RA target genes. Altogether, our data show that RA signaling rewires the epigenome and chromatin architecture to promote the expression of target genes during early embryonic development.

## RESULTS

### Dynamic binding of Retinoic Acid Receptor to chromatin during zebrafish development

To study the functions of the retinoic acid signaling pathway during zebrafish embryonic development, we first characterized by ChIPmentaton (ChIP-seq coupled to Tn5-mediated TAGmentation of chromatin) (40) in whole embryos the chromatin binding dynamics of the main RAR isoform in zebrafish, RARαa (41). For this, we selected three embryonic stages: 80% of epiboly (80epi, 8.3 hours post fertilization, hpf), which corresponds to gastrulation; 5 somite stage (5ss, 11.6 hpf), which corresponds to early neurulation and segmentation; and 24 hpf, which corresponds to the phylotypic stage. We obtained a total of 8,084 high confident RARαa binding sites (BSs) throughout the three embryonic stages (Figure 1A). A *k-*means clustering analysis of RARαa BSs revealed 6 major clusters, 3 of them showing a marked dynamic behavior: cluster 4 was composed of early RARαa BSs peaking at 80epi; cluster 5 was composed of intermediate RARαa BSs peaking at 5ss; and cluster 3 was composed of late RARαa BSs peaking at 24 hpf (Figure 1A). Motif enrichment analyses of dynamic BSs confirmed the prevalence of the RAR:RXR binding motif and other NR family motifs in all clusters (Figure 1B). In addition, early RARαa BSs showed also enrichment of the Sox family motif (Sox3), suggesting a cooperation between RARαa and the neuroectodermal TF Sox3 during early development that is consistent with the known role of RA inducing differentiation to neuroectoderm in ESCs (42). Early RARαa BSs also showed enrichment of the pluripotency TFs motif (Oct4-Sox2-Tcf-Nanog; Figure 1B), suggesting that RA signaling might have a role at pluripotency CREs.

**Figure 1.**
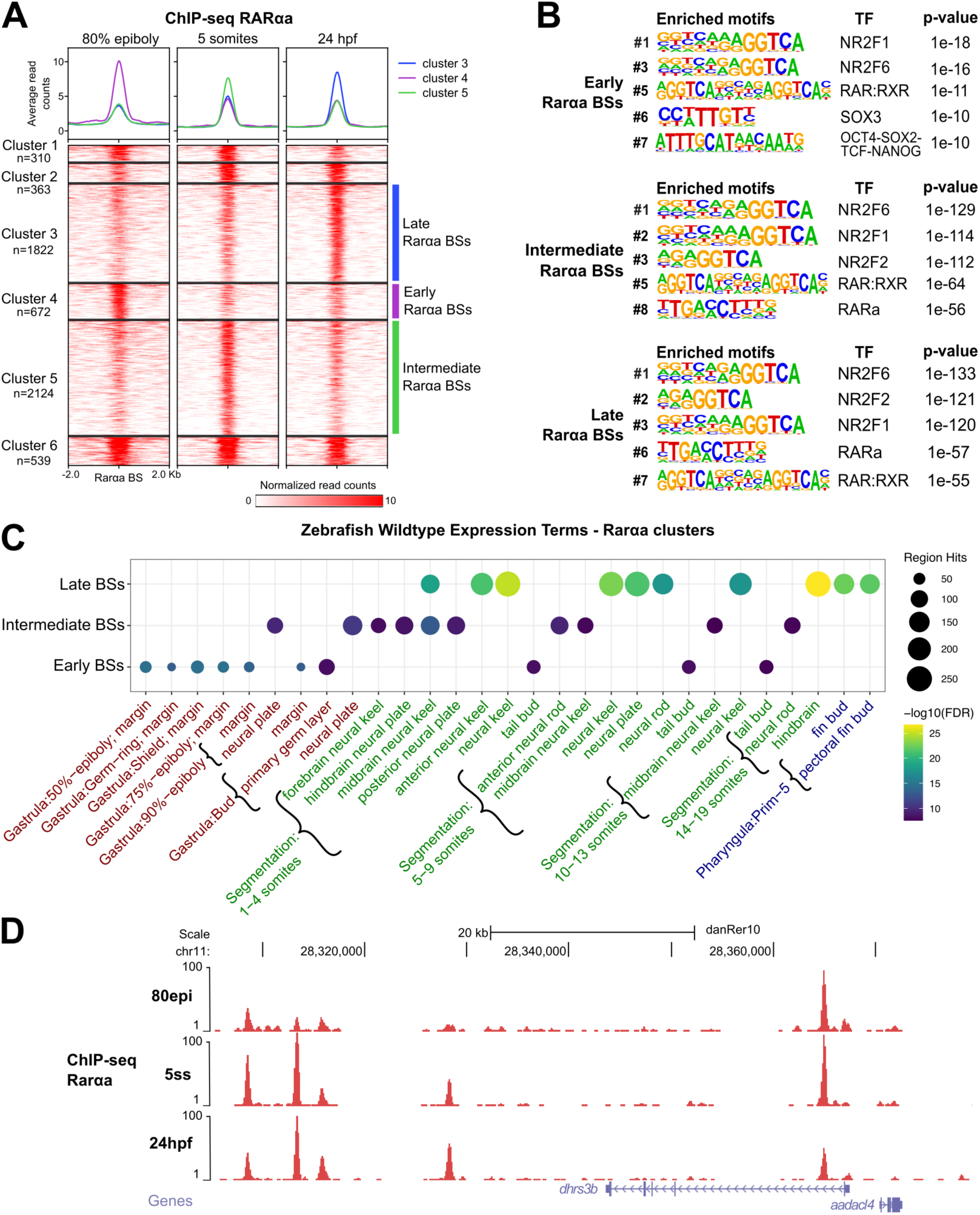
Dynamics of RARαa binding to chromatin during zebrafish embryonic development. **(A)** Heatmaps of the 8,084 RARαa BSs obtained from ChIP-seq in 80% of epiboly, 5 somites and 24hpf stages (n = 2 biological replicates per stage). Peaks were clustered using k-means clustering, obtaining six clusters, with three of them showing a clear dynamic behavior: cluster 4 (n = 672), or early BSs, cluster 5 (n = 2,059) or intermediate BSs, and cluster 3 (n = 1,764) or late BSs. Average profiles of clusters 4, 5 and 3 are shown on top. **(B)** Motif enrichment analysis of the early (top), intermediate (medium) and late (bottom) RARαa BSs. Five representative motifs of the top-10 were chosen. Motif logos are represented with their position in the top-10, the TF names and the enrichment p-values. **(C)** Zebrafish wild-type expression terms enriched for the genes associated with dynamic RARαa BSs. The top-10 terms for each stage have been combined. **(D)** Genome tracks of RARαa ChIP-seq at the indicated developmental stages showing signal intensities in the dhrs3b locus. The Genes track represents ENSEMBL annotated genes.

Next, we associated the dynamic RARαa BSs to their putative target genes using GREAT (see Methods) and found thousands of genes potentially regulated by RA signaling. Among them, there were many known RA targets in the three clusters, including RAR genes (*raraa*, *rarab*, *rarga* and *rargb*), RA metabolism genes (*aldh1a2*, *aldh1a3*, *dhrs3a*, *cyp26a1*, *cyp26b1*), early Hox genes (*hoxa1a*, *hoxb1a*, *hoxb1b*, *hoxc1a*, etc.), Meis genes (*meis1*, *meis2a* and *meis3*) and other reported RA targets (*fgf8a*, *pax6a/b, crabp2a*, *nrf1*) (Supplementary Dataset). Gene Ontology (GO) term enrichment analyses showed biological functions related to regulation of gene expression and embryonic patterning among the genes associated with the early RARαa BSs, development of the eye, digestive tract and brain for intermediate RARαa BSs and response to hormone for the late RARαa BSs (Suppl. Figure S1A), most of them corresponding to described functions of the RA signaling pathway and reflecting its high pleiotropy (1). Enrichment of genes belonging to the Notch signaling pathway was also found, illustrating the high interconnectivity among developmental signaling pathways in vertebrates (43). We also analyzed the enrichment in gene expression patterns using annotated information from the ZFIN database (see Methods). We found that genes associated with dynamic RARαa BSs exhibited a significant enrichment in gene expression primarily during the developmental stage in which the RARαa BSs reached maximum levels in each cluster (Figure 1C). In terms of anatomical structures, genes associated to early RARαa BSs were mostly expressed in the blastoderm margin, while genes associated to intermediate and late RARαa BSs were mostly expressed in the developing nervous system, including neural plate, neural keel, neural rod and hindbrain (Figure 1C). Expression in the pectoral fin buds also emerged in the cluster peaking at 24 hpf, which is consistent with the function of the RA pathway in the induction of the pectoral fin bud (44). Non-dynamic clusters of RARαa BSs were also associated to genes expressed in the developing nervous system during gastrulation, segmentation and pharyngula stages (data not shown). As an example, we found RARαa BSs with different temporal dynamics in close proximity to the *dhrs3b* gene, which encodes retinaldehyde reductase 3b, an enzyme that prevents the excessive accumulation of RA by the conversion of retinaldehyde back to retinol (Figure 1D) (45). Altogether, these data show that RARαa follows a dynamic chromatin binding behavior during zebrafish development, likely regulating genes that are related to known functions of the RA pathway, including but not restricted to the development of the nervous system and pectoral fin buds.

### Transcriptomic changes driven by RA treatment of zebrafish embryos

We aimed to identify the genes that are regulated by RA signaling during zebrafish development. For this purpose, we treated embryos with atRA at different timepoints and durations during development: gastrulation, by starting the treatment at 30% of epiboly and collecting embryos at 80epi; early segmentation and neurulation, by starting the treatment at 80epi and collecting embryos at 12 somites stage (12ss, 14 hpf); and the phylotypic stage, by starting the treatment also at 80epi and collecting embryos at 24 hpf (Figure 2A). AtRA treatment induced increased elongation of zebrafish embryos at the three tested conditions, but specially at 12ss, consistent with the role of RA signaling in body axis extension (Figure 2B).

**Figure 2.**
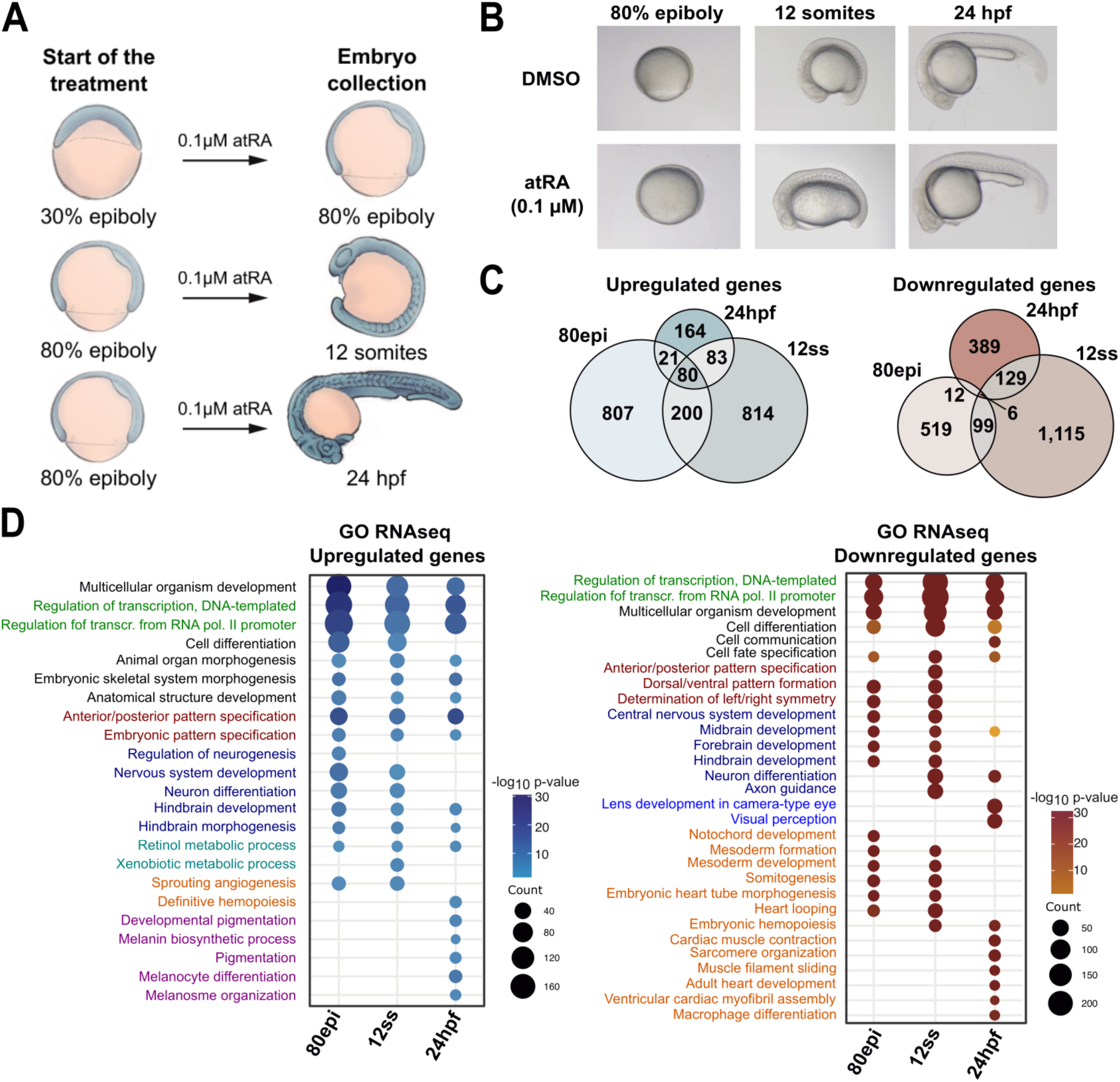
Transcriptomic effects of RA treatment to zebrafish embryos. **(A)** Picture depicting the treatments with 0.1 µM all-*trans* retinoic acid (atRA) of zebrafish embryos at different developmental stages. **(B)** Pictures of zebrafish embryos at 80% of epiboly, 12 somites and 24 hpf stages treated with DMSO or 0.1 µM atRA. **(C)** Venn diagrams showing the overlap between upregulated and downregulated genes at the analyzed stages. **(D)** GO Biological Process terms enriched for the upregulated and downregulated genes. The top-10 terms for each stage have been combined.

Next, we analyzed global gene expression changes induced by atRA treatment using RNA-seq in whole embryos. Differential expression analyses revealed 1,108, 1,177 and 348 upregulated, and 636, 1,349 and 536 downregulated genes at 80epi, 12ss and 24 hpf, respectively (FDR < 0.05, Fold-change ≥ 1.5; Suppl. Figure S1B). The lower number of differentially expressed genes (DEGs) at 24 hpf may be due to compensating mechanisms over developmental time, as the observed elongation phenotype is less evident at 24 hpf than at 12ss. We found a higher overlap among the upregulated genes at the three analyzed developmental stages, with 80 common genes, than among the downregulated genes, with only 6 common genes (Figure 2C). Common upregulated genes include well documented RA target genes, such as RA receptors (*raraa*, *rxrga* and *rxrgb*), genes involved in RA metabolism (*cyp26a1*, *dhrs3a* and *dhrs3b*), early Hox genes (*hoxa1a*, *hoxb1a*, *hoxb1b*, *hoxc1a*, etc) and Meis genes (*meis2b*, *meis3*), while common downregulated genes include *dmbx1a*, an homeobox gene involved in eye and tectum development (46) (Supplementary Dataset). GO term enrichment analyses of the DEGs showed an enrichment at the three stages of genes related to transcriptional regulation (i.e., TF genes), embryo patterning, neural development and differentiation for both up- and downregulated genes (Figure 2D), consistent with the known role of RA signaling in the development and patterning of the nervous system and with the regulation of downstream effector genes (1, 21). Pigmentation and melanocyte differentiation terms were also enriched in genes upregulated at 24 hpf, suggesting a role of RA signaling in the development of neural crest-derived cells, while terms related to mesoderm development, somitogenesis and heart development were enriched in the downregulated genes, suggesting that atRA treatment may have a negative effect in the specification of the mesoderm (Figure 2D). Consistently, gene expression patterns enriched for upregulated genes included rhombomeres of the hindbrain, spinal cord, somites and the neural crest, while those enriched for downregulated genes included also endoderm, otic vesicle and heart (Supplementary Fig. S1C). Altogether, these results highlight the importance of RA signaling regulation for multiple processes during early vertebrate development.

### RA treatment leads to an epigenome rewiring during early development

Previous studies showed changes in histone modifications and chromatin accessibility upon alteration of RA levels (32, 33). Thus, we wondered whether changes in gene expression induced by atRA treatment in zebrafish whole embryos could also occur together with alterations in CRE function. To address this, we performed ATAC-seq experiments in embryos treated with atRA as in Figure 2A. Statistical analyses of differentially accessible regions (DARs, FDR<0.1) showed 1,275 and 241 regions with increased and decreased accessibility, respectively, at 80epi (Figure 3A). However, only 53 and 44 peaks showed increased accessibility at 12ss and 24 hpf, respectively, and 1 and 7 peaks showed decreased accessibility at these stages (Figure 3A). Among the genes associated with increased DARs, we found RA pathway genes (*raraa*, *rxrga*, *roraa*, *dhrs3b*, *cyp26b1*), 3’ Hox genes (*hoxb1a*, *hoxb1b*, *hoxb2a*, *hoxd3a* and *hoxd4a*), Meis and Pbx genes (*meis2a*, *meis2b*, *pbx1b* and *pbx3b*) (Supplementary Dataset). GO enrichment analyses of genes associated with DARs showed biological functions consistent with our ChIP-seq and RNA-seq results, such as regulation of transcription, steroid hormone mediated signaling pathway, or nervous system development, even in the few peaks with increased accessibility at 12ss and 24 hpf (Figure 3B). Globally, there is a higher effect in chromatin accessibility of atRA treatment during gastrulation than during segmentation, suggesting that CREs responding to RA may become accessible during gastrulation, while treatment at later stages is not able to further increase their accessibility. The apparent discrepancy between ATAC-seq and RNA-seq data at 12ss and 24 hpf stages, with hundreds to thousands of DEGs but only few DARs, could be explained either by the fact that chromatin binding of RARαa and its targets may occur to already opened regions, without major alterations of global chromatin accessibility, although we cannot discard a higher dilution effect for the ATAC-seq in whole-embryo samples.

**Figure 3.**
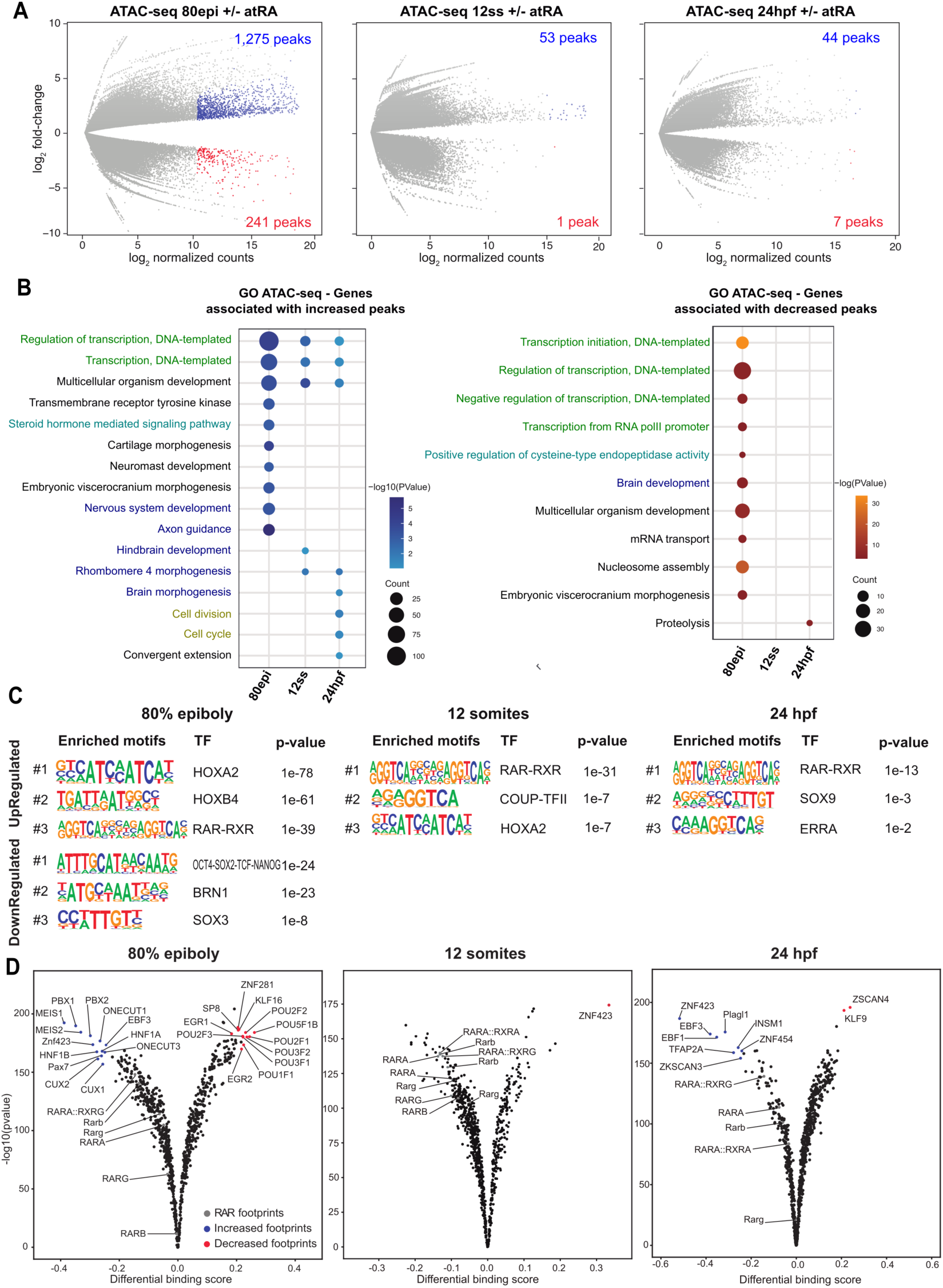
Changes in chromatin accessibility and TF binding induced by RA treatment. **(A)** Differential analyses of chromatin accessibility between atRA and DMSO treated embryos at 80% of epiboly, 12 somites and 24 hpf stages from ATAC-seq data (n = 2 biological replicates per stage and condition). The log_2_ p-value versus the log_2_ fold-change of accessibility is plotted. Regions showing statistically significant differential accessibility (adjusted P-value < 0.1) are highlighted in blue (increased) or red (decreased). The total number of differential peaks is shown inside the boxes. **(B)** GO Biological Process terms enriched for the genes associated with peaks with increased or decreased accessibility. The top-10 terms for each stage have been combined. **(C)** Motif enrichment analysis of the peaks with increased or decreased accessibility. The top-3 motifs have been selected for each stage. **(D)** Differential TF binding analysis in atRA and DMSO treated embryos at 80% of epiboly, 12 somites and 24 hpf stages from ATAC-seq data using TOBIAS. Volcano plots represent the differential binding score versus the -log_10_ p-value.

Motif enrichment analyses of DARs showed the RAR::RXR dimer binding motif among the top-3 motifs in regions with increased accessibility at the three embryonic stages (Figure 3C). We also found a high enrichment of HOX family motifs at 80epi, which was lower at 12ss and is consistent with Hox TFs being well-known targets of RA signaling (11). Finally, DARs with decreased accessibility upon atRA treatment at 80epi showed a high enrichment of pluripotency factor binding motifs, as well as SOX family motifs (Figure 3C), suggesting a function of RA signaling at CREs associated with pluripotency during early embryonic development. Indeed, we found binding of pluripotency TFs, including Pou5f3 (the zebrafish homolog of mammalian Oct4), Nanog and Sox2, specifically at DARs with decreased accessibility upon atRA treatment (Suppl. Figure S2A), and downregulation of *nanog*, *sox2* and *pou5f3* upon atRA treatment (Suppl. Figure S2B), which is in agreement with the reported repression of *pou5f1* by RAREs in zebrafish (47). These data suggest that RA signaling could be involved in the transition from pluripotent to differentiating cellular states.

Next, we aimed to detect TFs cooperating with RARαa in the response to RA signaling. For this, we performed footprinting analyses in our ATAC-seq data and calculated differential TF binding. Consistent with the differential accessibility analyses, we found significant changes in TF binding at 80epi but minor at 12ss and 24 hpf (Figure 3D). We detected increased binding of RAR at 80epi (8-10% increase over control, which is below our threshold of at least 15% change) and few changes at 12ss and 24 hpf. Interestingly, we found a higher chromatin binding of Meis and Pbx TFs at 80epi, which are known TFs cooperating with Hox proteins and have been recently described to cooperate with RA signaling in axial skeleton anterior-posterior patterning (48). Other TFs with increased footprint signal upon atRA treatment include Hnf1a, which is known to be activated by RA in the zebrafish posterior hindbrain (20); members of the cut-homeodomain family such as Cux2, which is a RA target that participates in the chicken limb positioning (49); Pax7, which is a marker of muscle stem cells whose expression is enhanced by RA (50); and Znf423, a TF involved in the midline patterning of the nervous system that is required for RA-induced differentiation (51). Surprisingly, the footprint of ZNF423 is increased by atRA treatment at 80epi but decreased at 12ss, coinciding with the alternate up- and downregulation of *znf423* gene at these stages, while at 24 hpf the footprint is increased again with no detected gene miss-expression, which suggests a complex regulation of this gene. Among the differentially bound TFs identified at 24 hpf, we detected increased footprints of the regulator of neural crest development Tfap2a (52), consistent with a possible role of RA signaling in neural crest-derived cells (Figure 2E). On the other hand, POU family of homeobox TFs showed decreased chromatin binding, consistent with motif analyses and chromatin closing of CREs related to pluripotency and with repression of *pou5f1* by RA (47), as well as Znf281, an inhibitor or RA-induced neuronal differentiation (53), Sp8, a regulator of Fgf8 and limb outgrowth (54), Klf16 and Egr1/2. Altogether, these data suggest that RA signaling promotes target gene expression during early development in cooperation with Hox/Meis/Pbx TFs and participates in the transition from pluripotency to lineage specification and differentiation.

### atRA treatment induces RARαa chromatin binding

AtRA treatment promotes chromatin accessibility at more than a thousand CREs detected by ATAC-seq during development (Figure 3). To assess what proportion of these CREs are opened directly or indirectly by increased RARαa binding, we first performed ChIP-seq of RARαa at 80epi embryos treated with atRA as described above (Figure 2A). Differential binding analysis found 986 peaks with increased RARαa binding and only 27 peaks with decreased binding upon atRA treatment (Figure 4A and Supplementary Fig. S3A), which validates *raraa* gene increased expression and confirms that exposure to RA at this stage is stimulating the binding of this RAR to chromatin. RARαa binding was highly specific, showing high enrichment of the RAR and NR family motifs in the peaks with increased RARαa binding (Suppl. Figure S3B). Genes associated to the new RARαa BSs in 80epi were enriched in expression in the neural keel and spinal cord during segmentation stages (Suppl. Figure S3C), suggesting that atRA treatment is causing a premature expression of RA target genes. Indeed, genes associated to BSs with increased RARαa binding were mostly upregulated (Figure 4B). This is illustrated by the *nr2f5* gene, which is upregulated by atRA treatment and has 5 RARαa BSs in proximity that arise upon RA stimulation (Figure 4C).

**Figure 4.**
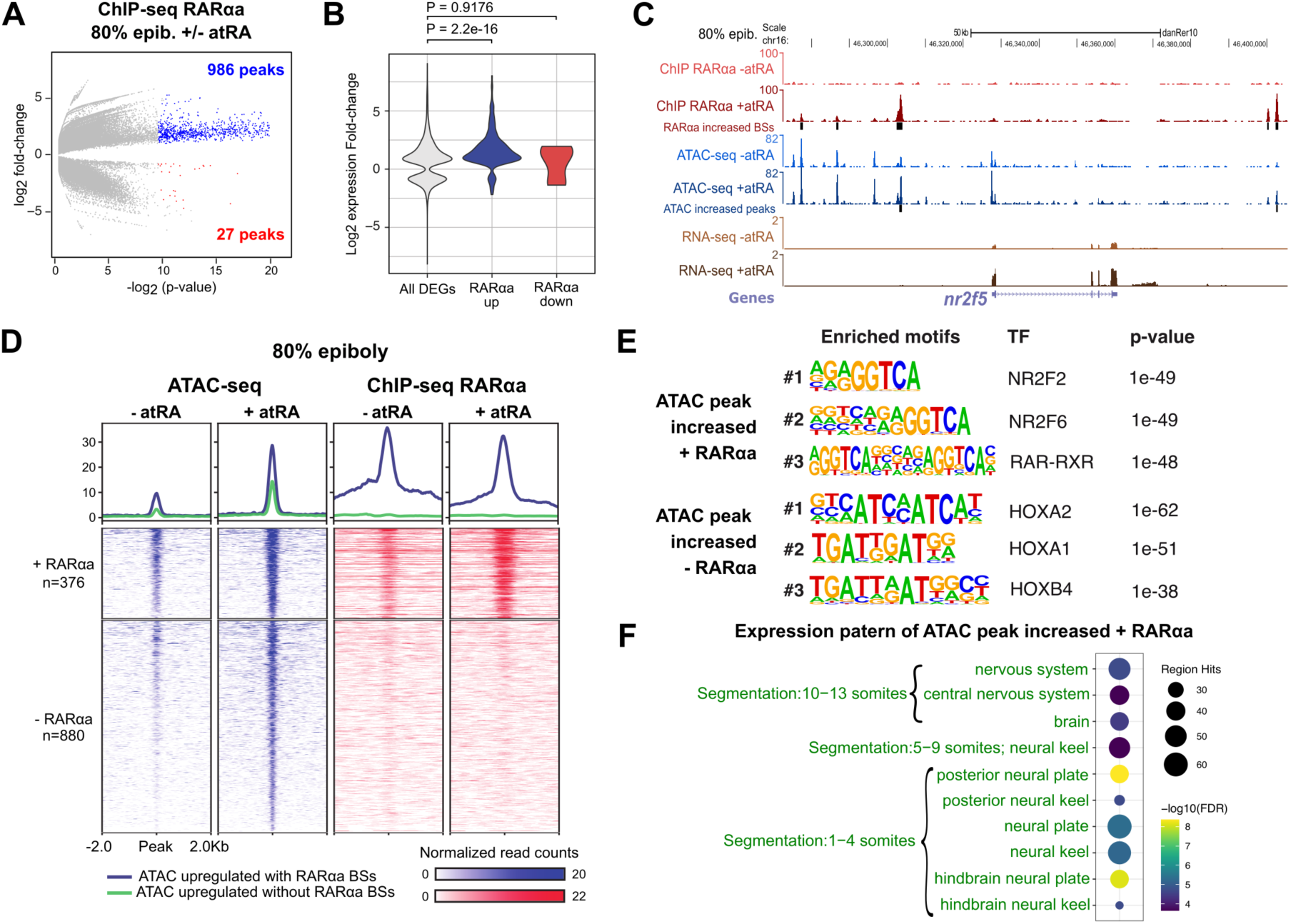
RA treatment leads to increased chromatin binding of its receptor RARαa. **(A)** Differential analysis of RARαa chromatin binding by ChIP-seq between atRA treated and control embryos at 80% of epiboly stage (n = 2 biological replicates per stage and condition). The log_2_ p-value versus the log_2_ fold-change of ChIP-seq signal are plotted. Regions showing statistically significant differential binding (adjusted P-value < 0.05) are highlighted in blue (increased) or red (decreased). The total number of differential peaks is shown inside the box. **(B)** Violin plots showing the distribution of log_2_ fold-change of expression (RNA-seq) all DEGs and those associated with increased or decreased chromatin binding of RARαa. **(C)** Genome tracks of RARαa ChIP-seq, ATAC-seq and RNA-seq at 80% of epiboly stage showing signal intensities in the *nr2f5* locus. The Genes track represents ENSEMBL annotated genes. **(D)** Heatmaps of the 1,275 ATAC-seq peaks with increased accessibility in atRA treated embryos at 80% of epiboly stage, separating those overlapping (n = 376) or not (n = 880) RARαa peaks. Average profiles of both groups are shown on top. **(E)** Motif enrichment analysis of the peaks with increased accessibility and RARαa binding or without RARαa. The top-3 motifs have been selected for each stage. **(F)** Zebrafish wildtype expression terms enriched for the genes associated with increased ATAC-seq peaks and RARαa binding. No enriched GO terms were found for the genes associated with increased ATAC-seq peaks but without RARαa binding.

Next, to distinguish between direct effects from atRA treatment that are carried out by RARαa from indirect effects caused by downstream regulators, we analyzed what proportion of the DARs with increased accessibility overlapped with RARαa binding at 80epi. We found that 376 out of 1275 (29.5 %) DARs with increased accessibility overlapped with RARαa BSs, which showed on average increased RARαa binding (Figure 4D), suggesting that almost one third of the effects of atRA on chromatin accessibility were due to a direct function of RARαa. In fact, these peaks showed a strong enrichment of the RAR and NR family binding motifs (Figure 4E). On the other hand, DARs with increased accessibility but no RARαa binding were highly enriched in HOX binding motifs (Figure 4E), suggesting that these TFs could be involved in the response to RA at the chromatin level and downstream of the RA receptor, which would be consistent with *Hox* genes being targets of RA signaling (24). GO enrichment analyses of genes associated to both groups of peaks showed an enrichment in genes expressed in the nervous system during segmentation stages for peaks with RARαa binding (Figure 4F), but no enrichment for peaks without RARαa binding. Taking together, our data indicate that epigenomic changes induced by atRA are due both to a direct function of RARαa in chromatin and to the indirect action of downstream effectors, among which Hox TFs could be involved.

### Chromatin binding of Hoxb1b, Meis2b and Sox3 TFs is stimulated by RA

We have shown so far that RA signaling promotes target gene expression by a rewiring of the chromatin accessibility landscape in which RAR and other downstream TFs may be involved, including Hox proteins. Indeed, we confirmed that multiple *hox* genes are upregulated upon atRA treatment and that multiple RARes near *hox* clusters show increased RARαa binding (Suppl. Figure S4). It is worth noting that most RARαa BSs responding to atRA treatment are located in the 3’ region of the *hox* clusters and that 3’ *hox* genes show a higher overexpression in response to atRA than 5’ *hox* genes (Suppl. Figure S4), consistent with the reported colinear activation of *hox* genes by RA signaling in chick embryos (28). Therefore, we wondered whether the transcriptional activation of 3’ *hox* genes by RA could result in increased chromatin binding of these TFs. To answer this, we performed ChIP-seq experiments at 80epi stage upon atRA treatment pulling down Hoxb1b, an anterior Hox TF involved in AP patterning whose expression is stimulated by atRA treatment. Differential binding analyses showed increased chromatin binding in 2,774 Hoxb1b BSs, while only 173 BSs showed decreased binding (Figure 5A and Supplementary Fig. S5A). In addition, since Meis and Pbx TFs are well-known co-factors of Hox proteins and given that we see the increased footprint signals of these TFs detected at 80epi (Figure 3D), we took advantage of the availability of the zebrafish Meis2b antibody and performed ChIP-seq experiments at 80epi upon atRA treatment. We detected 1,948 and 226 peaks with increased or decreased Meis2b binding, respectively (Figure 5A and Supplementary Fig. S5A). These results confirm that RA signaling promotes the chromatin binding of Hox and Meis TFs.

**Figure 5.**
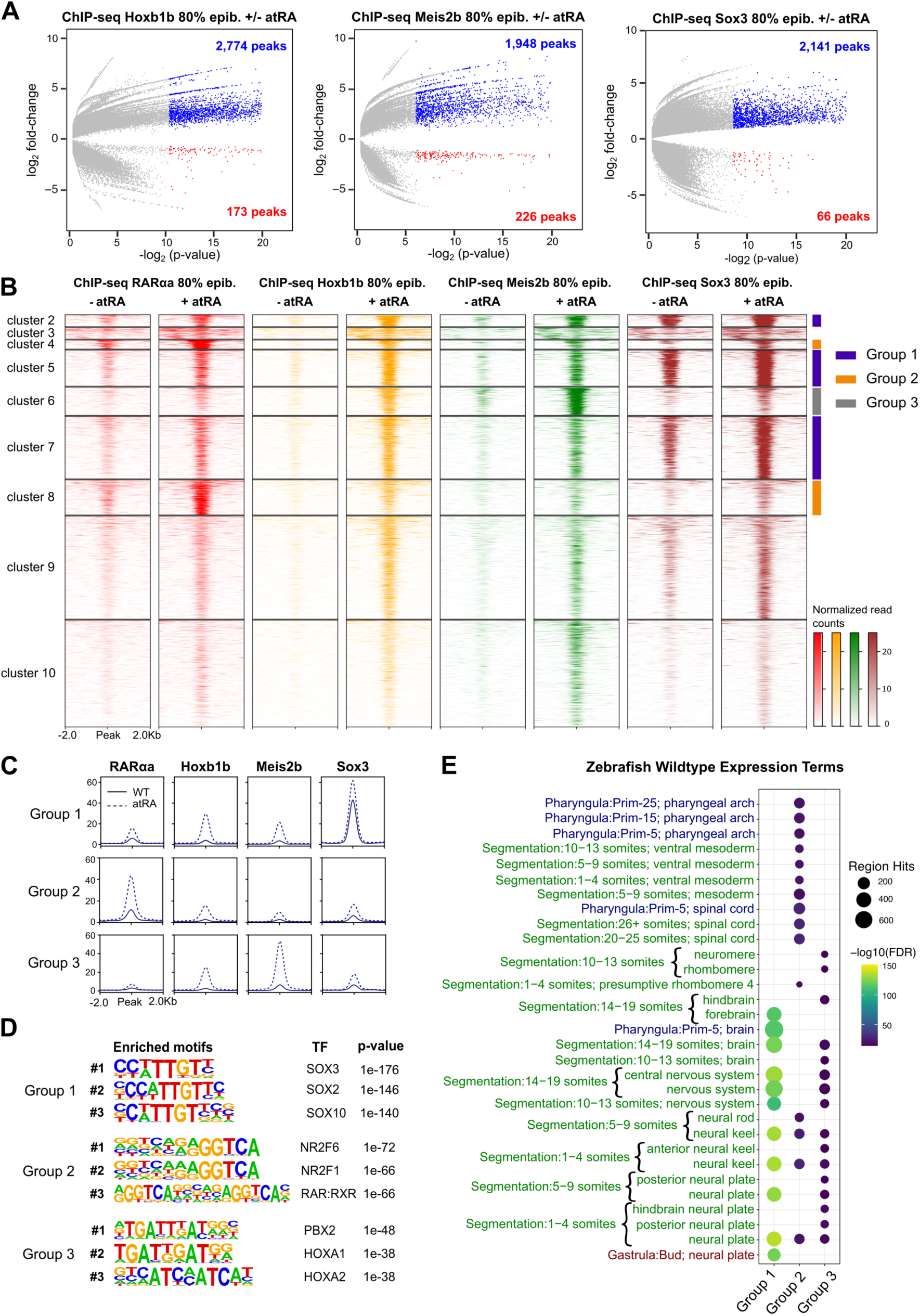
Chromatin binding of Hoxb1b, Meis2b and Sox3 is increased upon RA treatment. **(A)** Differential analysis of Hoxb1b, Meis2b and Sox3 chromatin binding by ChIP-seq between atRA treated and control embryos at 80% of epiboly stage (n = 2 biological replicates per condition). The log_2_ p-value versus the log_2_ fold-change of ChIP-seq signal are plotted. Regions showing statistically significant differential binding (adjusted P-value < 0.05) are highlighted in blue (increased) or red (decreased). The total number of differential peaks is shown inside the boxes. **(B)** Heatmaps of the combined 4,751 ChIP-seq peaks with increased binding of either RARαa, Hoxb1b, Meis2b or Sox3 obtained from ChIP-seq in 80% of epiboly. Peaks were clustered using *k*-means clustering, obtaining ten clusters, six of which were selected and combined in three groups with similar binding profiles of the four TFs: group 1 (n = 1,286, blue), group 2 (n = 515, orange) and group 3 (n = 328), grey. Cluster 1 was too small to plot and was omitted. **(C)** Average profiles of groups 1, 2 and 3 of ChIP-seq peaks with increased TF binding upon atRA treatment. **(D)** Motif enrichment analyses of the groups 1, 2 and 3 of ChIP-seq peaks with increased TF binding upon atRA treatment. The top-3 motifs have been selected for each group. **(E)** Zebrafish wild-type expression terms enriched for the genes associated with groups 1, 2 and 3 of ChIP-seq peaks with increased TF binding upon atRA treatment. The top-20 terms for each stage have been combined.

Motif enrichment analyses of the increased Hoxb1b and Meis2b BSs showed a strong enrichment of the HOX and PBX family motifs, consistent with the known cooperative role of Pbx and Meis TFs with Hox proteins (29), but also an enrichment of Sox family binding motifs (Supplementary Fig. S5B). The latter result, together with the enrichment of Sox motifs at early RARαa BSs (Figure 1B), suggests a cooperation among RA, Hox/Meis and Sox TFs. Thus, we decided to perform additional ChIP-seq experiments at 80epi with atRA treatment by pulling down the neuroectodermal TF Sox3. Differential binding analyses detected 2,141 and 66 Sox3 BSs with increased or decreased binding, respectively (Figure 5A and Supplementary Fig. S5A), with a high enrichment of Sox family motifs in the sites with increased binding (Supplementary Fig. S5B). Analyses of the expression pattern of genes associated to sites with increased binding of Hoxb1b, Meis2b or Sox3 showed a high overlap of expression in neural structures, including neural plate, neural keel, brain and spinal cord mainly during segmentation stages (Supplementary Fig. S5C). Altogether, these data indicate a cooperation of RA, Hox/Meis and Sox3 TFs during the early specification of the nervous system in zebrafish in response to RA signaling.

Next, we aimed to differentiate distinct functions of RARαa, Hoxb1b, Meis2b and Sox3 during early zebrafish development. For that purpose, we merged the TF BSs detected by ChIP-seq to be increased by atRA treatment and performed unsupervised *k*-means clustering. We detected 10 clusters with different TF profiles, selected those with the clearest differences in TF binding and grouped those with similar behaviors (Figure 5B). Thus, group 1 of BSs was composed of clusters 2, 5 and 7, and showed high levels of Sox3 occupancy that were further increased by atRA treatment and moderate increase in Hoxb1b and Meis2b binding; group 2 of BSs was composed of clusters 4 and 8, and showed high levels of RARαa upon atRA treatment, but low binding of the other TFs; and group 3 of BSs corresponded to cluster 6 and showed high occupancy of Meis2b upon atRA treatment and a moderate increase in Hoxb1b binding (Figure 5C). Motif enrichment analyses confirmed that group 1 corresponded mostly to Sox BSs, group 2 was composed of RAR BSs and group 3 were mainly Hox/Meis/Pbx BSs (Figure 5D). To distinguish specific functions of these groups of BSs, we associated them to their putative target genes and found that all of them were enriched in neural expression during segmentation stages, including neural plate, neural keel and brain (Figure 5E). However, there was a specific enrichment of expression in the spinal cord, the ventral mesoderm, and the pharyngeal arches slightly later in development (late segmentation and pharyngula stages) for the group 2, which corresponded to RARαa BSs (Figure 5E). Altogether, these data suggest that, at the analyzed stages of zebrafish development, RA signaling and its receptor RARαa cooperate with Hoxb1b/Meis2b and Sox3 in the gene regulatory network of the central nervous system early development, while it contributes to the development of the spinal cord, the ventral mesoderm and the pharyngeal arches independently of these TFs.

### RA signaling rewires promoter 3D interactions of target genes

We wondered whether atRA treatment could lead to the connection of CREs and genes regulated by RA signaling by chromatin 3D interactions. For this, we performed HiChIP experiments, which allow the analysis of chromatin 3D interactions at higher resolution than HiC by concentrating on those involving a protein or histone modification of interest. We analyzed whole embryos at 80epi stage, treated or not with atRA as in Figure 2A, by pulling down histone H3 lysine 4 trimethylation (H3K4me3), a mark of active promoters. Using this approach, we calculated differential promoter loops in both conditions, finding 931 and 958 loops with increased or reduced contacts, respectively, upon atRA treatment (Figure 6A). To see whether differential loops were associated with RA signaling, we calculated the overlap between loop anchors and RARαa BSs. Interestingly, we found that 53.6 % of loops with increased contacts upon atRA treatment contained RARαa BSs, versus 43 % of stable loops and 36.8 % of decreased loops (Figure 6B), a difference that was statistically significant (P < 0.00001). However, this was not the case of Hoxb1b, Meis2b and Sox3, since loop anchors with increased contacts were similarly or less occupied by these TFs than stable loops (Supplementary Figure S6A). These data suggest that RARαa-bound CREs engage into chromatin 3D contacts that are stimulated by RA signaling.

**Figure 6.**
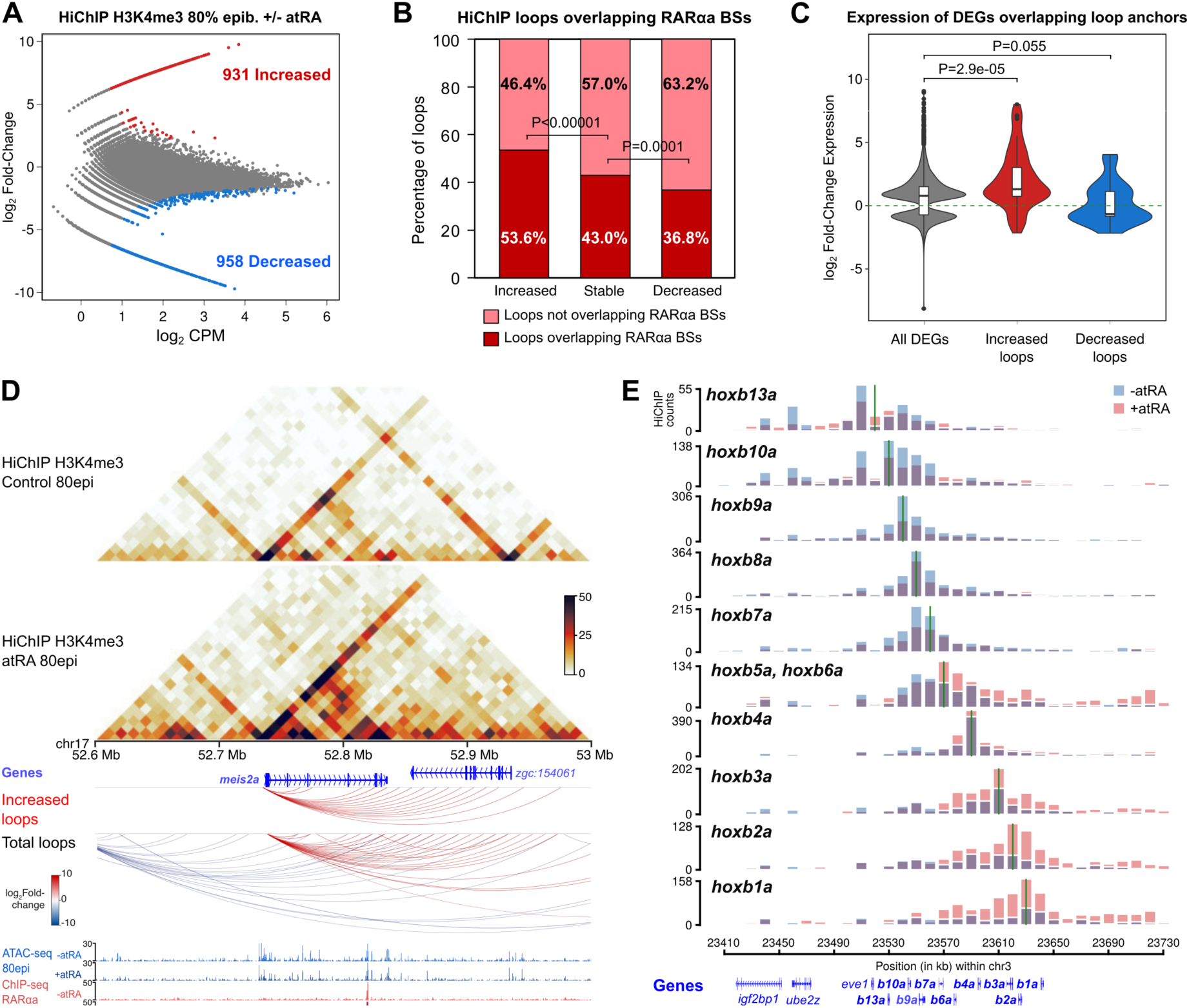
Rewiring of promoter 3D interactions by RA signaling. **(A)** Differential analyses of H3K4me3 HiChIP loops between control and atRA treated embryos at 80epi stage (n = 2 biological replicates per condition) at 10-Kb resolution. The log_2_ normalized counts per million (CPM) of control reads versus the log_2_ fold-change of contacts are plotted. Loops showing a statistically significant differential intensity (FDR < 0.05) are highlighted in red (increased) or blue (decreased). **(B)** Percentage of loops showing RARαa binding at least in one anchor for increased, stable and decreased loops. **(C)** Violing plots showing the expression fold-change in atRA treated embryos at 80epi of all DEGs and those associated with increased and decreased loops. **(D)** From top to bottom, heatmaps showing H3K4me3 HiChIP signal in control and atRA treated embryos, annotated genes, HiChIP loops increased by atRA treatment (FDR < 0.05), total HiChIP loops and tracks with ATAC-seq and RARαa ChIP-seq in control and atRA treated embryos at 80epi stage, in a 400-Kb region of chromosome 17 containing the upregulated gene *meis2a*. **(E)** Virtual 4C showing contact quantification calculated from H3K4me3 HiChIP data using as viewpoints 10-kb bins containing the gene promoters in the HoxBa cluster, both in control and atRA treated embryos at 80epi stage. A green vertical line indicates the bin used as viewpoint in each case. Annotated genes are shown at the bottom. Boxplots in C show: center line, median; box limits, upper and lower quartiles; whiskers, 1.5x interquartile range; notches, 95% confidence interval of the median. Statistical significance was assessed using a two-sided Wilcoxon’s rank sum test in C, and with a two-sided Fisher’s exact test in B.

Next, to test whether the changes in loop intensity were associated with changes in gene expression, we calculated the distribution of gene expression changes for DEGs overlapping increased or decreased loops. This analysis showed that RA target genes associated with increased promoter loops were significantly upregulated (P = 2.9e-5), while those associated with decreased promoter loops tended to be downregulated (P = 0.055; Figure 6C), indicating that changes in promoter 3D interactions often reflect changes in gene expression. Indeed, DEGs associated with increased loops were enriched in AP pattern specification, while those associated with decreased loops were enriched in somitogenesis and dorsal/ventral pattern formation, among other functions (Supplementary Figure S6B). These data suggest that RA signaling induces a rewiring of promoter 3D interactions of responsive genes that is concomitant with changes in their expression. The latter observation is illustrated by several RA target genes reported here. First, the *meis2a* gene is upregulated in response to atRA treatment, show near ATAC-seq and ChIP-seq peaks responding to atRA and its promoter is engaged in chromatin 3D interactions that are stimulated by RA (Figure 6D). On the other hand, the *pou5f3* gene is downregulated by atRA treatment and its promoter shows decreased contacts with nearby CREs (Supplementary Figure S6C).

Finally, we analyzed changes in chromatin 3D interactions at the HoxB cluster by calculating virtual 4C contacts from the HiChIP data for every HoxB gene promoter. Figure 6E shows that promoter interactions of 3’ HoxB genes, including *hoxb1a*, *hoxb2a*, *hoxb3a*, *hoxb4a*, *hoxb5a* and *hoxb6a*, are increased upon atRA treatment at the 3’ region of the cluster and the downstream gene desert. However, promoter interactions of 5’ HoxB genes, including *hoxb7a*, *hoxb8a*, *hoxb9a*, *hoxb10a* and *hoxb13a* show mainly decreased interactions upon atRA treatment with the 5’ region of the cluster and the upstream gene desert. Increased chromatin interactions at the downstream gene desert coincide with CREs showing enhanced accessibility and RARαa binding upon atRA treatment (Supplementary Figure S4D), as well as increased expression of the 3’ HoxB genes (Supplementary Figure S6D). These data support the idea that RA signaling promotes the expression of AP patterning genes by increased promoter 3D interactions with CREs activated by RARαa.

## DISCUSSION

In this work, we have used an integrative approach consisting of epigenomic, transcriptomic and chromatin conformation experiments, to get insight into the mechanisms by which RA signaling controls the expression of its target genes during early embryonic development. Using whole zebrafish embryos as a model, we leveraged TF ChIP-seq, ATAC-seq, RNA-seq and HiChIP experiments to show that this important signaling pathway rewires the embryo epigenome and reorganizes chromatin architecture, cooperating with downstream TFs to control its gene regulatory network.

Only few studies have addressed the genome-wide effects of RA signaling *in vivo*. In this sense, a recent report used dissected embryonic trunks from *Aldh1a2* knockout mice and identified CREs switching epigenomic state in the absence of RA near well-known RA target genes in the trunk (32). More recently, another study used endodermal cells from zebrafish embryos treated with RA or the inverse agonist BMS493, and found alterations in chromatin accessibility associated with genes involved in pancreatic development (33). Here, we have taken advantage of a zebrafish-specific antibody to profile the dynamic chromatin binding of the RA receptor RARαa by ChIP-seq through early development, covering from gastrulation to the phylotypic stage. Using this approach, we have identified dynamic RARαa BSs peaking at different stages (Figure 1) and have shown that early RARαa BSs, which are highly bound during gastrulation, are enriched in the DNA binding motif of the pluripotency TFs Oct4-Sox2-Tcf-Nanog. This is consistent with previous observations in embryonic carcinoma cells differentiating to primitive endoderm by RA treatment, in which RAR/RXR dimers bind to CREs occupied by pluripotency factors in undifferentiated cells, switching to Sox17 BSs in differentiated cells (55). However, RARαa BSs peaking later in development are highly enriched in RAR/RXR binding motifs and associated with genes expressed in the central nervous system and pectoral fins, suggesting that RARs may regulate the transition from pluripotency to patterning and differentiation processes.

Two main strategies have been used so far to identify RA functions and target genes in development: loss-of-function approaches, through the genetic knockout of RA producing enzymes (e.g. Aldh1a2) or the use of RA antagonists, and gain-of-function approaches, through the treatment with RA (56). Since zebrafish embryos can be treated with RA by just adding it to the embryo medium, we have taken advantage of this and treated them with 0.1 µM atRA at different embryonic stages, observing increased body axis elongation at all the analyzed stages (Figure 2). In contrast, previous studies using higher concentrations of atRA (1 µM) showed the opposite effect, i.e. body axis truncation (33), suggesting that the effect of RA signaling is highly dose-dependent. Indeed, lower concentrations of atRA require more exposure time to produce morphological effects (57). Using this approach, we have analyzed the transcriptomic changes driven by RA treatment at different developmental stages (Figure 2) and have observed that RA leads to a miss-regulation of more genes at the earlier stages, including gastrulation and early neurulation and segmentation, while treatment until 24 hpf provokes milder effects. Genes up-regulated at all stages include bona-fide RA target genes, such as *RAR* genes, RA metabolism genes, anterior *Hox* genes and *Meis* genes, as well as other genes involved in anterior/posterior pattern specification and nervous system development. These observations are consistent with the reported role of RA signaling in the patterning of the central nervous system and the pectoral fin bud during zebrafish pre-segmentation stages (44). Interestingly, genes stimulated by RA at later stages show enrichment of melanocyte development and neural crest-expressing genes, suggesting a function of RA signaling in neural crest development that agrees with its reported role in the generation and migration of neural crest cells in both avian and mammalian models (58, 59, 60).

We have also analyzed here the effects of RA treatment on chromatin accessibility by ATAC-seq (Figure 3). We have found more than a thousand putative CREs that gain accessibility upon atRA treatment during gastrulation, a third of which correspond to RARαa BSs and the remaining ones being enriched in Hox motifs (Figure 4). In contrast, putative CREs losing accessibility are occupied by pluripotency TFs, including Sox2, Nanog and Pou5f3 (the zebrafish ortholog of Oct4) (Supplementary Figure S2). These genes are also down-regulated at this stage, consistent with a possible role of RA signaling in exiting from pluripotent states and with a previous report showing reduced expression of pluripotency genes in the RA-induced differentiation to neural and endoderm fates (61). Surprisingly and in contrast to changes at the transcriptomic level, very few CREs show RA-induced changes in chromatin accessibility at later stages. This discrepancy may be due to CREs responding to RA being rewired at earlier stages, during gastrulation, and/or to RARαa binding to already accessible chromatin that is not opened further. Indeed, the latter possibility has been previously observed for the glucocorticoid receptor (62), and a recent study in MCF-7 cells showed that either RA or TGF-ß treatment induced both concordant and discordant changes in accessibility and gene expression (63). We cannot discard though a higher dilution effect on ATAC-seq data at later developmental stages due to the whole-embryo approach.

Estimation of differential TF binding upon RA treatment using footprint analyses confirms that Pou family of TFs show a reduced chromatin binding at early stages and that there is an increased binding of Meis and Pbx families. This is consistent with the enrichment of the Hox binding motif at CREs with increased accessibility, since these are common Hox co-factors (29). Anterior Hox genes are indeed well-known targets of RA signaling (24, 25, 26), and we show using TF ChIP-seq during gastrulation that not only RARαa binding is stimulated by RA (Figure 4), but also Hoxb1b, the earliest HoxB gene to be expressed, and Meis2b also increase their chromatin binding (Figure 5). Moreover, we also report the involvement of the early neuroectodermal TF Sox3, whose chromatin binding is also stimulated by RA. This observation agrees with a previous report showing that *SOX3* expression is stimulated by RAR/RXR binding elements (64), but little is known about this interaction and our observations provide a new branch of RA signaling downstream of RARs to promote neural development. Indeed, we show that RARαa, Hoxb1b, Meis2b and Sox3 regulate genes required for development of the central nervous system. In contrast, RARαa participates in additional functions independently of those TFs, such as the development of the ventral mesoderm and the pharyngeal arches, probably in cooperation with other developmental regulators not identified here. This illustrates the pleiotropy and multifunctional character of the RA signaling pathway during embryonic development.

Finally, we have investigated the molecular mechanisms connecting RA-induced transcriptomic and enhancer rewiring. Enhancers usually interact with their target promoters by chromatin 3D interactions that are believed to be essential for target gene expression (34). We show here using HiChIP that RA treatment of zebrafish embryos leads to a rewiring of chromatin architecture, with almost two thousand promoter interactions showing increased or decreased contacts (Figure 6). Promoter loops that change intensity by RA signaling are connected to genes whose transcriptional response to RA goes in the same direction, indicating a consistent connection between enhancer-promoter interactions and target gene expression. This has been previously shown for other differentiation processes *in vitro*, including neural and erythroid differentiation (37, 38). Regarding TF binding, we detect a higher enrichment of RARαa BSs at the anchors of loops induced by RA, suggesting a connection between TF binding and target gene expression by the establishment of chromatin 3D interactions. However, Hoxb1b, Meis2b and Sox3 do not show this enrichment, which could be due to their implication in pre-stablished loops or to a lower detection of differential loops in which they participate. In any case, the existence of chromatin loops altered by RA and connected to RA-responding genes is illustrated by bona-fide induced and repressed targets, such as *meis2a* and *pou5f3*, respectively. Further evidence comes from the regulation of the HoxB cluster by RA, for which we show that the anterior HoxB genes are engaged in increased chromatin interactions within the cluster and with the 3’ gene desert, while posterior HoxB genes show decreased interactions. These observations are consistent with the specific upregulation of anterior HoxB genes and with the existence of RARαa BSs only in the 3’ region and gene desert of the cluster (Supplementary Figure S4) and confirms the importance of RA signaling in regulating the establishment of the anterior/posterior body axis. Consistently, a previous study in ESCs showed increased interactions between enhancers containing RAREs and *Hoxa1* gene that promoted its expression (39), while here we show that this is a general effect concerning anterior Hox genes and the main RA target genes.

## CONCLUSIONS

We show here that RA signaling leads to changes in gene expression and chromatin accessibility during embryonic development. It promotes chromatin binding of RARαa and the downstream TFs Hoxb1b, Meis2b and Sox3 to promote the anterior/posterior patterning and development of the central nervous system and other structures. Furthermore, we connect changes in TF binding and target gene expression by showing that RA signaling rewires chromatin 3D interactions, providing a molecular mechanism by which developmental signaling pathways control the expression of their target genes. Further studies will be required to see whether this mechanism also applies to other signaling pathways or is more prevalent for some of them, including RA.

## METHODS

### Animal experimentation and embryo treatments

Wild-type AB/Tübingen zebrafish strains were maintained and bred under standard conditions. All experiments involving animals conform national and European Community standards for the use of animals in experimentation and were approved by the Ethical Committees from the University Pablo de Olavide, CSIC and the Andalusian government. Embryos were treated with 0.1 µM atRA or DMSO and collected at the indicated developmental stages (Figure 2A).

### RNA-seq

For total RNA extraction, atRA or DMSO treated embryos were collected, de-chorionated with 30 mg/mL Pronase (Roche) and suspended in TRIsure (Bioline). 30 embryos were used for 80epi stage, 20 embryos for 12ss and 15 embryos for 24hpf. RNA was then purified with Direct-zol RNA miniprep kit (Zymo Research) and treated with TURBO DNA free kit (Invitrogen). Three biological replicates were used for each analyzed treatment and stage. Illumina libraries were constructed and sequenced in a BGISEQ-500 single-end lane producing around 50 million (M) of 50-bp reads. Reads were aligned to the GRCz10 (danRer10) zebrafish genome assembly using STAR 2.5.3a (65) and counted using the htseq-count tool from the HTSeq 0.8.0 toolkit (66). Differential gene expression analysis was performed using the DESeq2 1.18.1 package in R 3.4.3 (67), setting a corrected P value < 0.05 and fold-change > 1.5 as the cutoff for statistical significance of the differential expression. Enrichment of GO Biological Process terms was calculated using David 6.8 (68), with a false discovery rate (FDR)-corrected P value < 0.05 as statistical cutoff. Enrichment of zebrafish wild-type expression patterns was calculated using previously published code (69).

### ATAC-seq

ATAC-seq assays were performed using standard protocols (70, 71), with minor modifications. Briefly, atRA and DMSO treated embryos were de-chorionated with 30 mg/mL Pronase (Roche). 30 embryos were used for 80epi stage, 10 embryos for 12ss and 5 embryos for 24hpf. Yolk was dissolved with Ginzburg Ring Finger (55 mM NaCl, 1.8 mM KCl, 1.15 mM NaHCO_3_) by pipetting and shaking 5 min at 1100 rpm. Deyolked embryos were collected by centrifugation for 5 min at 500g 4°C. Supernatant was removed and embryos washed with PBS. Then, embryos were lysed in 50 µl of Lysis Buffer (10 mM Tris-HCl pH 7.4, 10 mM NaCl, 3 mM MgCl_2_, 0.1% NP-40) by pipetting up and down. From the whole cell lysate, 80,000 cells were used for TAGmentation, which were centrifuged for 10 min at 500g 4°C and resuspended in 50 µl of the Transposition Reaction, containing 1.25 µl of Tn5 enzyme and TAGmentation Buffer (10 mM Tris-HCl pH 8.0, 5 mM MgCl2, 10 % w/v dimethylformamide), and incubated for 30 min at 37°C. Immediately after TAGmentation, DNA was purified using the MinElute PCR Purification Kit (Qiagen) and eluted in 10 µl. Libraries were generated by PCR amplification using NEBNext High-Fidelity 2X PCR Master Mix (NEB). The resulting libraries were purified using MinElute PCR Purification Kit (Qiagen), multiplexed and sequenced in a HiSeq 4000 pair-end lane producing around 100M of 49-bp pair end reads per sample.

### ChIP-seq by ChIPmentation

ChIP-seq of RARαa, Hoxb1b, Meis2b and Sox3 were performed by ChIPmentation, which incorporates Tn5-mediated TAGmentation of immunoprecipitated DNA, as previously described (40, 69). Briefly, 400 zebrafish atRA treated or control embryos were dechorionated with 300 µg/ml pronase, fixed for 10 min in 1% paraformaldehyde (in 200 mM phosphate buffer) at room temperature, quenched for 5 min with 0.125 M glycine, washed in PBS and frozen at -80°C. Fixed embryos were homogenized in 2 ml cell lysis buffer (10 mM Tris-HCl pH 7.5, 10 mM NaCl, 0.3% NP-40, 1x Roche Complete protease inhibitors cocktail) with a Dounce Homogenizer on ice and centrifuged 5 min 2,300g at 4°C. Pelleted nuclei were resuspended in 333 µl of nuclear lysis buffer (50 mM Tris-HCl pH 7.5, 10 mM EDTA, 1% SDS, 1x Roche Complete protease inhibitors cocktail), kept 5 min on ice and diluted with 667 µl of ChIP dilution buffer (16.7 mM Tris-HCl pH 7.5, 1.2 mM EDTA, 167 mM NaCl, 0.01% SDS, 1.1% Triton-X100). Then, chromatin was sonicated in a Covaris M220 sonicator (duty cycle 10%, PIP 75W, 100 cycles/burst, 10 min) and centrifuged 5 min 18,000g at 4°C. The recovered supernatant, which contained soluble chromatin, was used for ChIP or frozen at -80°C after checking the size of the sonicated chromatin. Four 250 µl aliquots of sonicated chromatin were used for each independent ChIP experiment, and each aliquot incubated with 2 µg of antibody (anti-Rarαa GTX124492, anti-Hoxb1b GTX128322, anti-Meis2b GTX127229, or anti-Sox3 GTX132494) and rotated overnight at 4°C. Next day, 20 µl of protein G Dynabeads (Invitrogen) per aliquot were washed twice with ChIP dilution buffer and resuspended in 50 µl/aliquot of the same solution. Immunoprecipitated chromatin was then incubated with washed beads for 1 hour rotating at 4°C and washed twice sequentially with wash buffer 1 (20 mM Tris-HCl pH 7.5, 2 mM EDTA, 150 mM NaCl, 1% SDS, 1% Triton-X100), wash buffer 2 (20 mM Tris-HCl pH 7.5, 2 mM EDTA, 500 mM NaCl, 0.1% SDS, 1% Triton-X100), wash buffer 3 (10 mM Tris-HCl pH 7.5, 1 mM EDTA, 250 mM LiCl, 1% NP-40, 1% Na-deoxycholate) and 10 mM Tris-HCl pH 8.0, using a cold magnet (Invitrogen). Then, beads were resuspended in 25 µl of TAGmentation reaction mix (10 mM Tris-HCl pH 8.0, 5 mM MgCl2, 10 % w/v dimethylformamide), added 1 µl of Tn5 enzyme and incubated 1 min at 37°C. TAGmentation reaction was put in the cold magnet and the supernatant discarded. Beads were washed twice again with wash buffer 1 and 1x TE and eluted twice for 15 min in 100 µl of elution buffer (50 mM NaHCO3 pH 8.8, 1% SDS). The 200 µl of eluted chromatin per aliquot were then decrosslinked by adding 10 µl of 4M NaCl and 1 µl of 10 mg/ml proteinase K and incubating at 65°C for 6 hours. DNA was purified using Minelute PCR Purification Kit (Qiagen), pooling all aliquots in a single column, and eluted in 20 µl. Library preparation was performed as previously described for ATAC-seq (see above). Libraries were multiplexed and sequenced in HiSeq 4000 or DNBseq pair-end lanes producing around 20M of 49-bp or 50-bp paired-end reads per sample, respectively.

### ChIP-seq and ATAC-seq data analyses

ChIP-seq and ATAC-seq reads were aligned to the GRCz10 (danRer10) zebrafish genome assembly using Bowtie2 2.3.5 (72) and those pairs separated by more than 2 Kb were removed. For ATAC-seq, the Tn5 cutting site was determined as the position -4 (minus strand) or +5 (plus strand) from each read start, and this position was extended 5 bp in both directions. Conversion of SAM alignment files to BAM was performed using Samtools 1.9 (73). Conversion of BAM to BED files, and peak analyses, such as overlaps or merges, were carried out using the Bedtools 2.29.2 suite (74). Conversion of BED to BigWig files was performed using the genomecov tool from Bedtools and the wigToBigWig utility from UCSC (75). For ATAC-seq, peaks were called using MACS2 2.1.1.20160309 algorithm (76) with an FDR < 0.05 for each replicate and merged in a single pool of peaks that was used to calculate differentially accessible sites with DESeq2 1.18.1 package in R 3.4.3 (67), setting a corrected P value < 0.1 as the cutoff for statistical significance of the differential accessibility. For ChIP-seq, RARαa peaks with a global IDR < 0.01 were called using the IDR framework (idr 0.1 version) to obtain high confidence peaks based on replicate information, as previously described (77), and these peaks were used for clustering analyses. Alternatively, RARαa, Hoxb1b, Meis2b and Sox3 peaks with an FDR < 0.05 were called with MACS2 and used for differential binding calculation with DESeq2 1.18.1 package in R 3.4.3 (67), setting a corrected P value < 0.05 as the cutoff for statistical significance of the differential binding. For visualization purposes, reads were extended 100 bp for ATAC-seq and 300 bp for ChIP-seq. For data comparison, all ChIP-seq and ATAC-seq experiments used were normalized using reads falling into peaks to counteract differences in background levels between experiments and replicates (69).

Heatmaps and average profiles of ChIP-seq and ATAC-seq data were generated using computeMatrix, plotHeatmap and plotProfile tools from the Deeptools 3.5 toolkit (78). TF motif enrichment was calculated using the script FindMotifsGenome.pl from Homer 4.11 software (79), with standard parameters. For gene assignment to ChIP and ATAC peaks, coordinates were converted to Zv9 (danRer7) genome using the Liftover tool of the UCSC Genome Browser (75) and assigned to genes using the GREAT 3.0.0 tool (80), with the basal plus extension association rule with standard parameters (5 Kb upstream, 1 Kb downstream, 1 Mb maximum extension). This tool was also used to calculate Gene Ontology and zebrafish wildtype expression term enrichment with standard parameters (significant by both region-based binomial and gene-based hypergeometric test, FDR < 0.05). Peak *k*-means clustering was calculated using seqMiner (81). For footprinting analyses, we used TOBIAS 0.12.9 (82). First, we performed bias correction using ATACorrect and calculated footprint scores with ScoreBigwig, both with standard parameters. Then, we used BINDetect to determine the differential TF binding for all vertebrate motifs in the JASPAR database (83). We considered as differentially bound those motifs with a linear fold-change ≥ 15% between atRA and DMSO treated embryos.

### HiChIP

HiChIP assays were performed as previously described (84). Briefly, 1,000 atRA-treated or control zebrafish embryos at 80% epiboly stage were dechorionated with 300 µg/ml pronase and transferred to 1 ml of Ginzburg fish ringer buffer (55 mM NaCl, 1.8 mM KCl, 1.25 mM NaHCO3). Yolks were disrupted by pipetting and shaking for 5 min at 1100 rpm. Embryos were then spinned-down and fixed as indicated above for ChIPmentation. Fixed embryos were homogenized in 5 ml cell lysis buffer (see above) with a Dounce Homogenizer on ice. Complete cell lysis generating nuclei was checked at the microscope with methylgreen-pyronin staining. Nuclei were then centrifuged 5 min at 600g at 4°C and in situ contact generation was performed as described (85), with modifications. For chromatin digestion, 8 µl of 50 U/µl DpnII restriction enzyme were used. Ligation was incubated overnight at 16°C shaking at 900 rpm. Both digestion and ligation efficiencies were monitored by taking control aliquots that were de-crosslinked, phenol-chloroform purified and loaded in an 0.7% agarose gel. The controls consisted of 5-µl aliquots before and after digestion (undigested and digested controls), a 10-µl aliquot before end repair that was ligated with overhang ends (3C control) and a 25-µl aliquot after ligation (ligation control).

After ligation, nuclei were pelleted at 2500g for 5 min at RT, resuspended in 495 µl of nuclear lysis buffer (see above) and kept 5 min on ice to lysate nuclei. Then, 1,980 µl of ChIP dilution buffer (see above) were added and sample split in three 1-ml aliquots that were sonicated in a Covaris M220 sonicator (duty cycle 10%, PIP 75W, 100 cycles/burst, 5 min). Then, sonicated chromatin was centrifuged for 15 min at 16,000g at 4°C and the supernatant transferred to a new tube. Sonication efficiency was checked using a 20-µl aliquot that was RNase A-treated, de-crosslinked, phenol-chloroform purified and loaded in a 0.7% agarose gel. After this, chromatin was pre-cleared with Dynabeads protein G (Invitrogen) rotating for 1 hour at 4°C, recovered to new tubes using a magnet and incubated overnight rotating at 4°C with 6.7 µg (20 µg total) of anti-H3K4me3 (Abcam ab8580) antibody per sample. Immunoprecipitated chromatin was then washed and eluted from beads as described (85). Before DNA purification, the three samples were mixed and split in two samples to generate later two independent libraries, increasing likelihood of library amplification over primer artifact amplification.

Biotin capture with Streptavidin C-1 beads (Invitrogen) and TAGmentation were performed as described (85). For library preparation, samples were put in a magnet, supernatant discarded, and beads resuspended in a 50-µl PCR mix containing 1x NEBNext High-Fidelity PCR Master Mix (NEB) and 0.5 µM of Nextera Ad1_noMX and Ad2.X primers. PCR was run for 5 cycles and then samples put in a magnet to separate beads. Then, cycle number for library preparation was estimated by qPCR taking a 2-µl aliquot from the samples, and the remaining PCR was run for the empirically determined number of cycles. Finally, libraries were recovered from beads using a magnet, pooled together, and purified using DNA Clean and Concentrator columns (Zymo Research), eluting in 20 µl of 10 mM Tris-HCl pH 8.0. Libraries were quantified in a Qubit machine and sequenced using DNBseq technology to generate around 500M of 50-bp paired-end reads.

### HiChIP data analyses

HiChIP paired-end reads were aligned to GRCz10 (danRer10) zebrafish genome assembly using the TADbit pipeline (86). Default settings were used to remove duplicate reads, assign reads to DpnII restriction fragments, filter for valid interactions, and generate binned interaction matrices with a 10-kb resolution. Data was visualized using the WashU Epigenome Browser (87) and the fancplot tool from the FAN-C 0.9.14 toolkit 2 (88). Since HiC normalization methods are not suitable for HiChIP data given the inherent scarcity of HiChIP matrices, we scaled the samples to the same number of valid read pairs.

For differential analysis of HiChIP loops between atRA-treated and control embryos we used the DiffAnalysisHiChIP tool of FitHiChIP 9.0 (89) with the following parameters: interaction type peak to all, bin size 10 Kb, distance threshold between 20 Kb and 20 Mb, FDR < 0.01, loose background for contact probability estimation [FitHiChIP(L)], coverage bias regression and merging of redundant loops. We considered a stringent set of loops consisting of the merge of those detected in both biological replicates per condition, with a differential FDR threshold of 0.05 and a fold-change threshold of 1.5. To avoid calling loops as differential due to different ChIP-seq coverage, we only considered loops involving H3K4me3 peaks not called as differential by EdgeR (i.e., categories ND-ND, LD-LD and ND-LD from the differential analysis output). Virtual 4C tracks of HoxB genes were generated from HiChIP interaction matrices. First, virtual 4C baits were determined by overlapping of HiChIP 10-kb bins with HoxB genes coordinates within chromosome 3. Then, we focused on a 320-kb locus around the HoxB cluster (chr3:23410000-23730000) and extracted all interaction counts from each single bait belonging to such locus.

### Statistical analyses

For comparison of data distribution, two-tailed Wilcoxon’s rank sum tests or Student’s *t*-tests were used. Statistical significance of contingency tables was assessed using the Fisher’s exact test.

## DECLARATIONS

### Ethics approval and consent to participate

All experiments involving animals conform national and European Community standards for the use of animals in experimentation and were approved by the Ethical Committees from the University Pablo de Olavide, CSIC and the Andalusian government.

### Consent for publication

Not applicable.

### Availability of data and materials

The RNA-seq, ATAC-seq, ChIP-seq and HiChIP data generated in this study have been deposited in the Gene Expression Omnibus (GEO) database under accession code GSE233698. The ChIP-seq data used in Supplementary Figure S2 can be accessed using codes GSE34684 and GSE39780.

## Funding

JJT was funded by the Spanish Ministerio de Ciencia e Innovación (PID2019-103921GB-I00) and the institutional grant Unidad de Excelencia María de Maeztu to CABD (CEX2020-001088-M). JMS-P was funded by Junta de Andalucía (ProyExcel_00363) and a postdoctoral fellowship (DOC_00512). PM-G was funded by a postdoctoral fellowship from Junta de Andalucía (DOC_00397).

### Author contributions

LG-F, MM-O, JLG-S and JMS-P conceived and designed the project; LG-F, MM-O, SN and SJ-G performed the experiments; LG-F, MM-O, PMM-G, JJT and JMS-P analyzed the data; LG-F, MM-O and JMS-P wrote the original manuscript. All authors contributed to the final manuscript.

## Acknowledgements

We dedicate this study to the memory of our friend, mentor and colleague, José Luis Gómez-Skarmeta and thank C. Paliou and M. Franke for critical reading of the manuscript.

