## Supplementary Figures for "Rewiring of the epigenome and chromatin architecture by retinoic acid signaling during zebrafish embryonic development"

[1] Centro Andaluz de Biología del Desarrollo (CABD), Consejo Superior de Investigaciones Científicas/Universidad Pablo de Olavide, 41013 Seville, Spain.

SUPPLEMENTARY MATERIAL

- Supplementary Figures S1 to S6

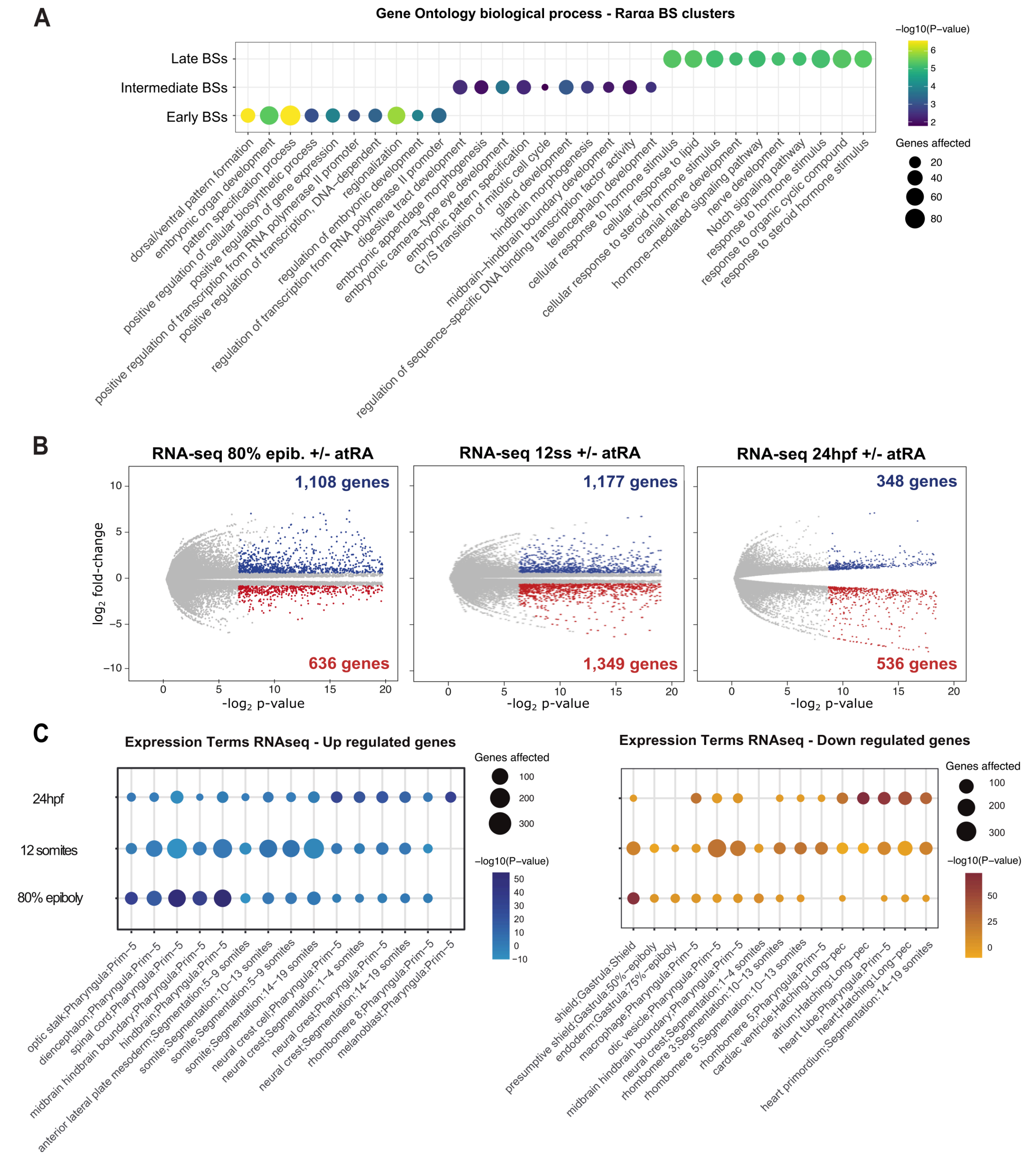


**Figure S1. (A)** GO Biological Process terms enriched for the genes associated with dynamic RARαa BSs. The top-10 terms for each stage have been combined. **(B)** Differential analyses of gene expression between atRA and DMSO treated embryos at 80% of epiboly, 12 somites and 24 hpf stages from RNA-seq data (n = 3 biological replicates per condition). The $\log_2$ p-value versus the $\log_2$ fold-change of expression is plotted. Transcripts showing a statistically significant differential expression (adjusted P-value < 0.05 and absolute fold-change > 1.5) are highlighted in blue (upregulated) or red (downregulated). The number of genes that correspond to the upregulated and downregulated transcripts are shown inside the boxes. **(C)** Zebrafish wild-type expression terms enriched for the upregulated and downregulated genes. The top-10 terms for each stage have been combined.

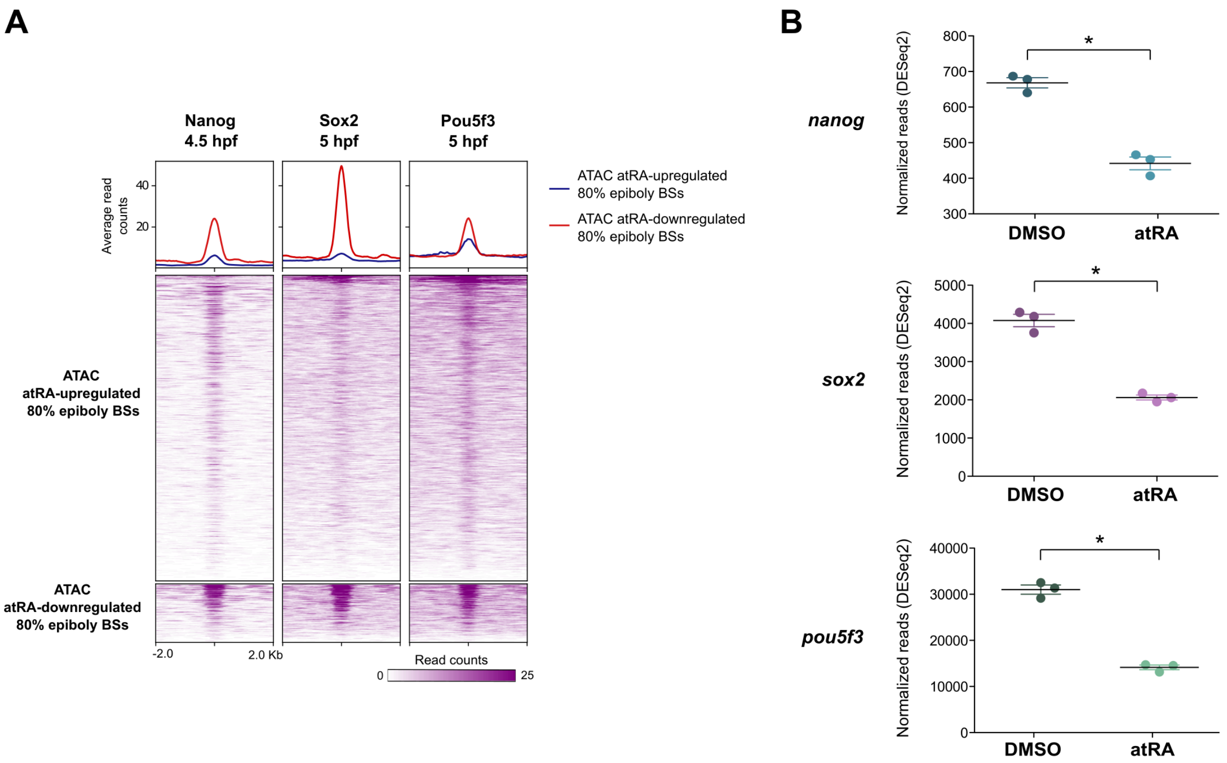


**Figure S2. (A)** Heatmaps showing ChIP-seq signal of Nanog (GSE34684), Sox2 (GSE39780) and Pou5f3 (GSE39780) at 4.5-5 hpf stages over the 1,275 ATAC-seq peaks with increased accessibility and the 241 ATAC-seq peaks with decreased accessibility in atRA treated embryos at 80% of epiboly stage. Average profiles are shown on top. **(B)** Dot plots showing expression levels of *nanog*, *sox2* and *pou5f3* genes in atRA and DMSO treated embryos at 80% of epiboly stage. Normalized read counts are plotted.

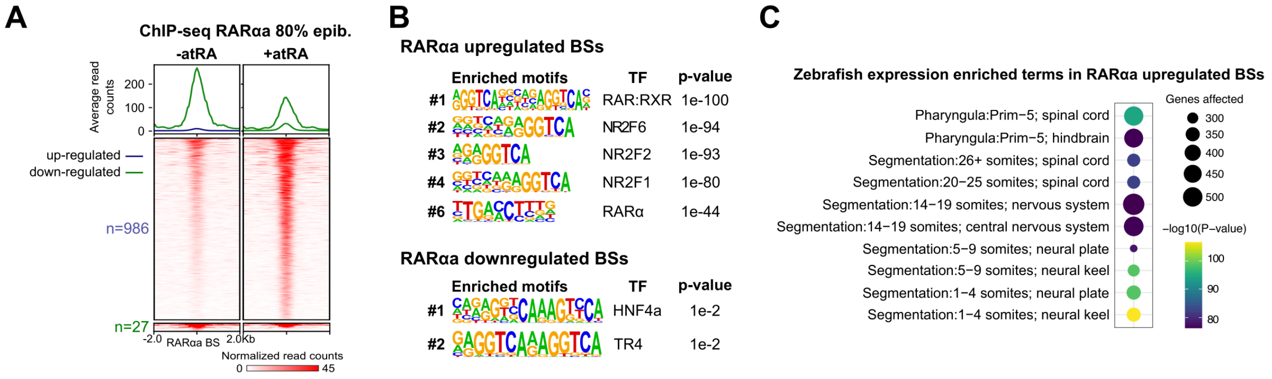


**Figure S3. (A)** Heatmaps showing RARαa ChIP-seq signal for the 986 and 27 peaks with increased or decreased RARαa binding, respectively. Average profiles of both groups are shown on top. **(B)** Motif enrichment analyses of the ChIP-seq peaks with increased or decreased RARαa binding. **(C)** Zebrafish wild-type expression terms enriched for the genes associated with ChIP-seq peaks with increased RARαa binding. No enriched terms were detected for the ChIP-seq peaks with decreased RARαa binding.

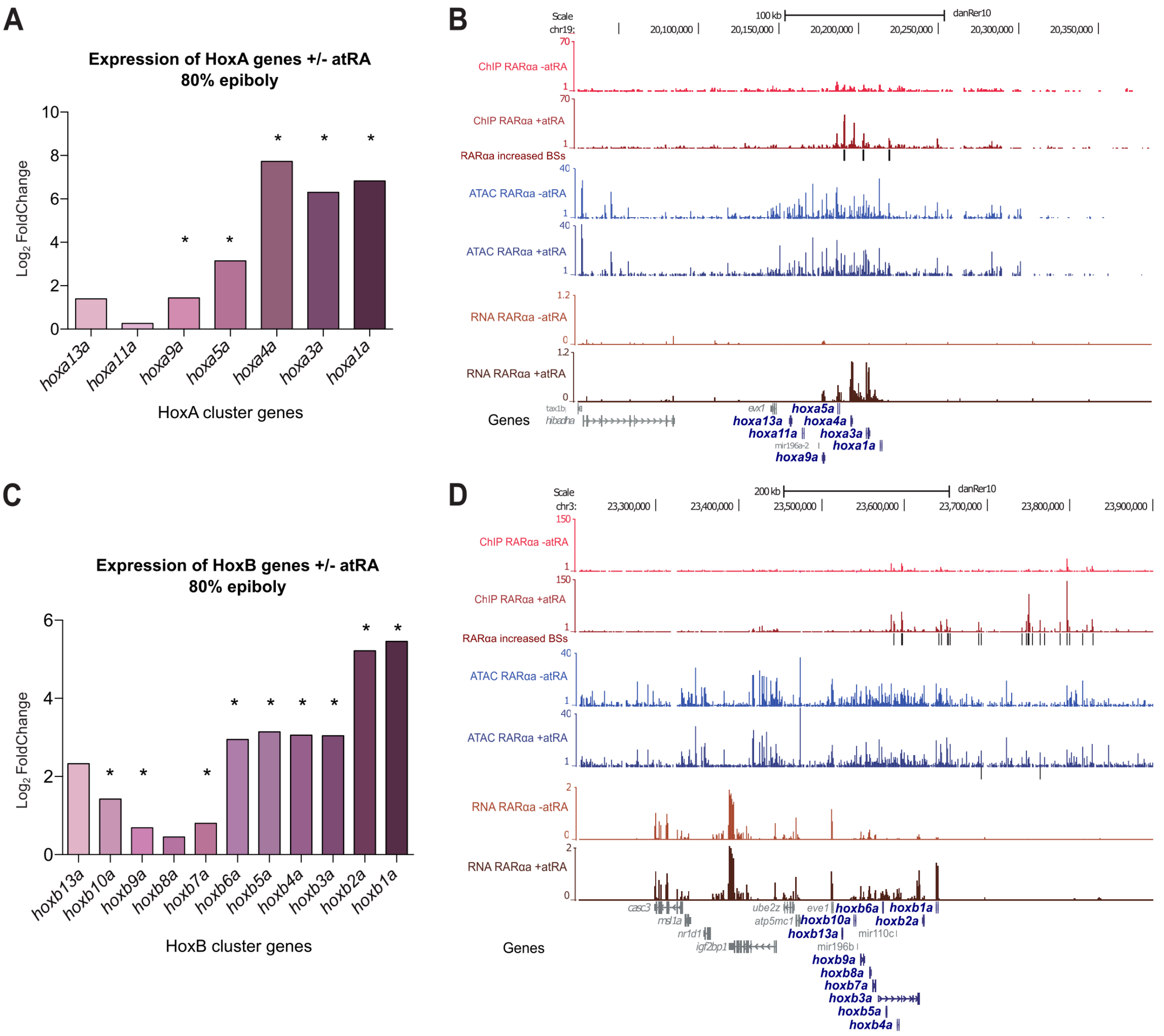


**Figure S4. (A)** Change in expression upon atRA treatment at 80% of epiboly stage embryos for the Hox genes in one of the HoxA clusters in zebrafish (HoxAa). The $\log_2$ fold-change of expression is plotted. **(B)** Genome tracks of RARαa ChIP-seq, ATAC-seq and RNA-seq at 80% of epiboly stage showing signal intensities in the HoxAa cluster. **(C)** Change in expression upon atRA treatment at 80% of epiboly stage embryos for the Hox genes in one of the HoxB clusters in zebrafish (HoxBa). The $\log_2$ fold-change of expression is plotted. **(D)** Genome tracks of RARαa ChIP-seq, ATAC-seq and RNA-seq at 80% of epiboly stage showing signal intensities in the HoxBa cluster. The Genes track represents ENSEMBL annotated genes. *, Adjusted P-value < 0.05 and absolute fold change > 1.5.

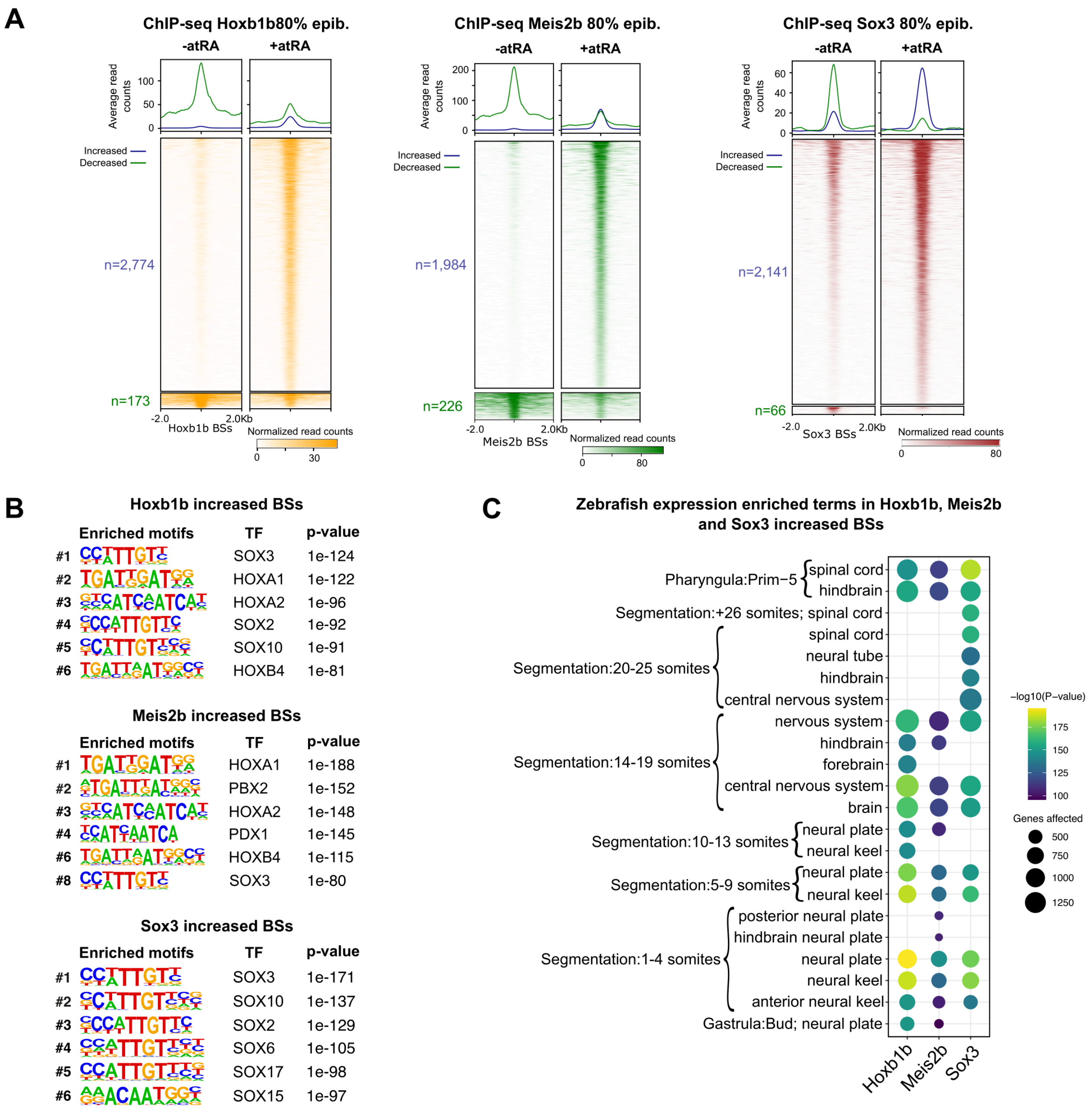


**Figure S5. (A)** Heatmaps showing Hoxb1b, Meis2b or Sox3 ChIP-seq signals for the peaks with increased or decreased binding of each TF. Average profiles are shown on top. **(B)** Motif enrichment analyses of the ChIP-seq peaks with increased or decreased Hoxb1b, Meis2b or Sox3 binding. Six representative motifs from the top-10 have been selected. **(C)** Zebrafish wild-type expression terms enriched for the genes associated with ChIP-seq peaks with increased binding of Hoxb1b, Meis2b or Sox3. The top-20 terms for each TF have been combined.

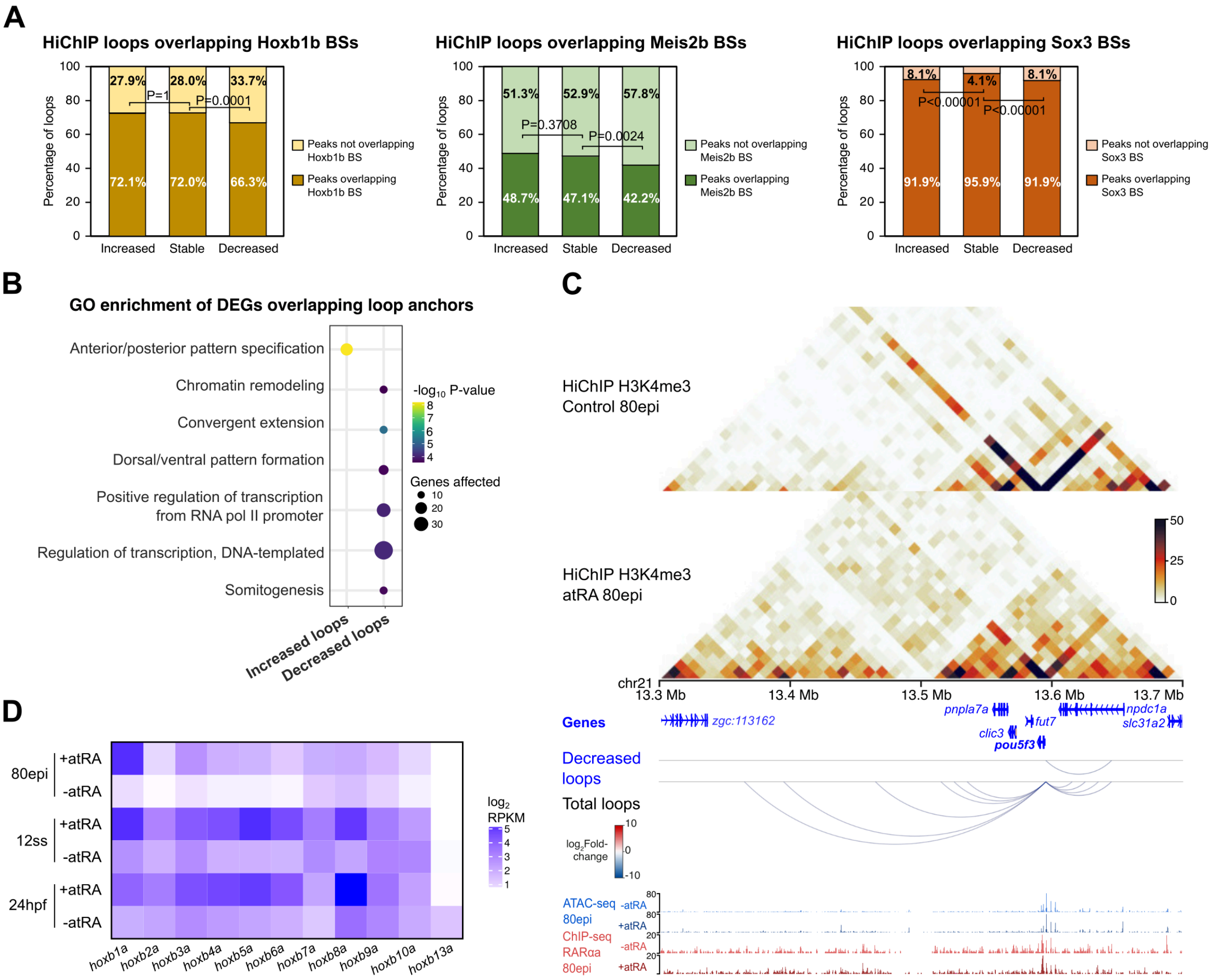


**Figure S6. (A)** Percentage of loops showing Hoxb1b (left), Meis2b (middle) and Sox3 (right) binding at least in one anchor for increased, stable and decreased H3K4me3 HiChIP loops upon atRA treatment. **(B)** GO Biological Process terms enriched for the differentially expressed genes associated with increased and decreased HiChIP loops. **(C)** From top to bottom, heatmaps showing H3K4me3 HiChIP signal in control and atRA treated embryos, annotated genes, HiChIP loops decreased by atRA treatment (FDR < 0.05), total HiChIP loops and tracks with ATAC-seq and RARαa ChIP-seq in control and atRA treated embryos at 80epi stage, in a 400-Kb region of chromosome 21 containing the downregulated gene *pou5f3*. **(D)** Heatmaps showing the normalized expression of genes in the HoxBa cluster in control and atRA treated embryos at 80epi, 12ss and 24hpf stages.
